# Sensorimotor transformation underlying odor-modulated locomotion in walking *Drosophila*

**DOI:** 10.1101/2022.03.15.484478

**Authors:** Liangyu Tao, Samuel P Wechsler, Vikas Bhandawat

## Abstract

Most real-world behaviors are performed with incomplete information. Odor-guided locomotion, an ecologically important behavior essential to an animal’s survival, is an example of such a behavior. Different odors activate different patterns of olfactory receptor neuron (ORN) classes providing information about which odor is present but does not provide any navigational information. In this study, we investigate the sensorimotor transformation that relates ORN activation to locomotion changes in *Drosophila* by optogenetically activating different combinations of ORN classes and measuring the resulting changes in locomotion. Three features describe this sensorimotor transformation: First, locomotion depends on both the instantaneous firing frequency (f) and its change (*df*); the two together serve as short-term memory that allows the fly to automatically adapt its motor program to sensory context. Second, the mapping between *f-df* and locomotor parameters such as speed or curvature is distinct for each pattern of activated ORNs. Finally, the sensorimotor mapping changes with time after odor exposure allowing integration of information over a longer timescale.

## Introduction

Locating ecologically important resources such as food and mates is critical to the survival of all animals (Lessing and Carlson, 1999). Odors emanating from these resources are an important source of information for locating and assessing them. Animals can smell odors from a long distance, but because at long distances there are no gradients (Murlis et al., 1992), and therefore little navigational information, finding the source of odor requires animals to act on incomplete information. Smelling an odor will often result in explorative search rather than navigational movements (Cardé and Willis, 2008; Murlis et al., 1992). During explorative search, at each instant the animal chooses between a vast number of possible actions; each action does not result in a defined outcome but might yield new information or close the door to this information. Many real-world behaviors have a similar structure and require continuous decision-making over extended periods with incomplete information (Yoo et al., 2021). The neural mechanisms underlying such behaviors are poorly understood; odor-guided locomotion provides a great opportunity to further understand this important class of behaviors.

Odor-gated exploratory search results in varied and distance-dependent effect on locomotion. At long distances from the source, odors stimulate locomotion and can direct locomotion in an upwind direction; these and other changes in locomotion often bring the animal closer to the odor source (Alvarez-Salvado et al., 2018; Budick and Dickinson, 2006; Wallraff, 2004; Weissburg, 2000; Willis and Arbas, 1998; Willis and Avondet, 2005; Wolf and Wehner, 2000). Closer to the odor source, decrease in walking speed, increased turning and other changes keeps the animal close to the resource. Overall animals use a suite of locomotor mechanisms to locate odor sources, home in on it, and interact and utilize resources important for survival.

Relating odor-driven changes in locomotion to stimulus encounter is a challenging problem because of two reasons both related to stimulus control: First, because of the nature of odor transport, the spatial and temporal pattern of odor experienced by the animal is varied and changes with environmental conditions and distance from the odor source making the characterization of odor stimulus itself difficult (Capelli et al., 2013; Celani et al., 2014; Elkinton et al., 1984; Murlis et al., 1992; Riffell et al., 2008). Second, how a given odor is encoded by olfactory neurons is dependent not just on the identity of the odor but also on odor concentration: Odors are detected by olfactory receptor neurons (ORNs) which each express one to few receptors that determine the ORN’s response profile and therefore forms an ORN class. The complement of ORN classes activated by a given odor depends on the odor concentration (Hallem and Carlson, 2006).

The first challenge of defining the odor stimulus experienced by the animal was addressed in works in moths when it became possible to record odor stimulus in flying moths (Mafra-Neto and Cardé, 1994; Vickers and Baker, 1994). Similar work has been performed in other animals including other insects (Dekker and Cardé, 2011; Thiery and Visser, 1986; van Breugel and Dickinson, 2014) and in mice and reveal many conserved mechanisms at play across the animal kingdom. These experiments represent significant progress in our understanding. However, two limitations remain. One limitation is relating these mechanisms to a model for locomotion. Attempts to model behavior have suggested that these mechanisms work well when the environment is predictable (Adden et al., 2020) but may not work well in an unpredictable environment (Willis and Arbas, 1998). Another limitation is that few studies have connected behavior to neural response because of the problem of replicating odor stimulus in an electrophysiological rig. These limitations mean that neural algorithms underlying odor-modulated locomotion remains poorly understood.

Another limitation of the work described above is that experiments were performed on a single odor or odor blend, therefore, the second challenge of relating the complement of ORNs activated by different odors and behavior remained unaddressed. Relating activation of different ORN classes to the movement of freely locomoting animals has predominantly been addressed only in *Drosophila* because of the genetic tools and the relative ease with which experiments can be performed on many flies. Much of this work has focused on the question of valence or what makes an odor attractive or repulsive (Badel et al., 2016; Bell and Wilson, 2016; Knaden et al., 2012; Tumkaya et al., 2021). To probe the mechanism of attraction – i.e., what are the changes in locomotion underlying attraction to odor in flies - we created a ring arena whose center had a fixed odor concentration, and the periphery did not have any odor (Jung et al., 2015).

Flies only experienced odors when they went inside the central odor-zone, therefore both the timing and concentration of odor was known. This study showed that different ORN classes activate many different motor parameters independently (Jung et al., 2015). This behavioral algorithm of ORNs modulating many different motor parameters independently can be executed by parallel pathways that connect each ORN class to multiple higher-order neurons: ORNs of a given class project to a single glomerulus in the antennal lobe. Each glomerulus is innervated by multiple types of second-order neurons called projection neurons (PNs) that include both excitatory and inhibitory uniglomerular PNs and multiglomerular PNs implying multiple parallel pathways downstream of each ORN class (Bates et al., 2020; Scheffer et al., 2020). Each PN contacts multiple third-order neurons providing even more opportunities for parallel computations (Schlegel et al., 2021).

Although the previous study showed that different ORNs affected motor parameters independently, much of the analysis was based on averages over minutes and did not provide any insight into how moment-by-moment firing of ORNs affects a fly’s locomotion; neither did it connect changes in motor parameters to changes in the distribution of the fly in the arena. Equally importantly, locomotion itself would affect sensory experience because the fly’s locomotion regulates the sampling dynamics of odor: if the fly darts in and out of the odor quickly, the responses to subsequent pulses of odors will be affected by the ongoing response to the first odor encounter. Sampling dynamics play an important role in mammalian odor coding (Wachowiak, 2011).

In this study, we address three unsolved problems. First, we obtain a moment-by-moment record of both the sensory and behavioral information by recording from ORNs and measuring changes in locomotion. Second, we create a generative model of locomotion to show that the measured changes indeed describe the fly’s overall behavior. Finally, we systematically activate multiple patterns of ORN classes to unravel the logic between patterns of active ORN and the resulting change in behavior. We solve the above problems by optogenetically activating different ORN classes and measuring behavioral changes in *Drosophila*. We also measure the ORN response to the stimulus experienced by the fly. This experimental design provides an accurate estimate of both the temporal pattern of activity in the ORNs as well as the identity of the ORN activated and allows us to accurately characterize the underlying sensorimotor transformation. We discover that the fly automatically adapts its locomotor strategy on both short and long timescales, and that its behavioral response depends on the complement of ORNs activated.

## Results

### Changes in the distribution of the fly depend on the combination of active ORN classes

We focused on subsets of ORN classes activated by a powerful attractant, apple cider vinegar. Apple cider vinegar activates seven ORN classes (Jung et al., 2015). We activated five of these ORN classes either singly or in combinations of two or three ORN classes using genetic lines that express the transcription factor *Gal4* under the control of olfactory receptor promoters (Ai et al., 2010; Couto et al., 2005; Fishilevich and Vosshall, 2005; Silbering et al., 2011); these genetic lines express the transcription factor *Gal4* in a single ORN class. *Gal4* was used to drive the expression of *CsChrimson* (*Chrimson* for short), a red light activated channel(Klapoetke et al., 2014). Flies that express *Chrimson* under the control of known ORN classes were placed in a small circular arena (8 cm in diameter) whose central region – a circular region 2 cm in diameter- had a fixed intensity of the light (Tao et al., 2020) (Figure 1A). As the fly walked into the region with the red light (light-zone or stimulated region), specific sets of ORNs were activated by the red light, allowing an assessment of the resulting behavioral change. Because flies’ photoreceptors have low sensitivity in the long wavelength, their behavioral response to red light itself is small. Chrimson requires retinal for its response to light and is fed to the flies by raising them on food containing retinal; flies raised on non-retinal food served as controls. As in previous studies (Jung et al., 2015; Tao et al., 2020), we first measured the fly’s baseline behavior for a three-minute period during which the light in the central area was off, turned the light on, and measured its behavior for an additional three minutes.

**Figure 1.**
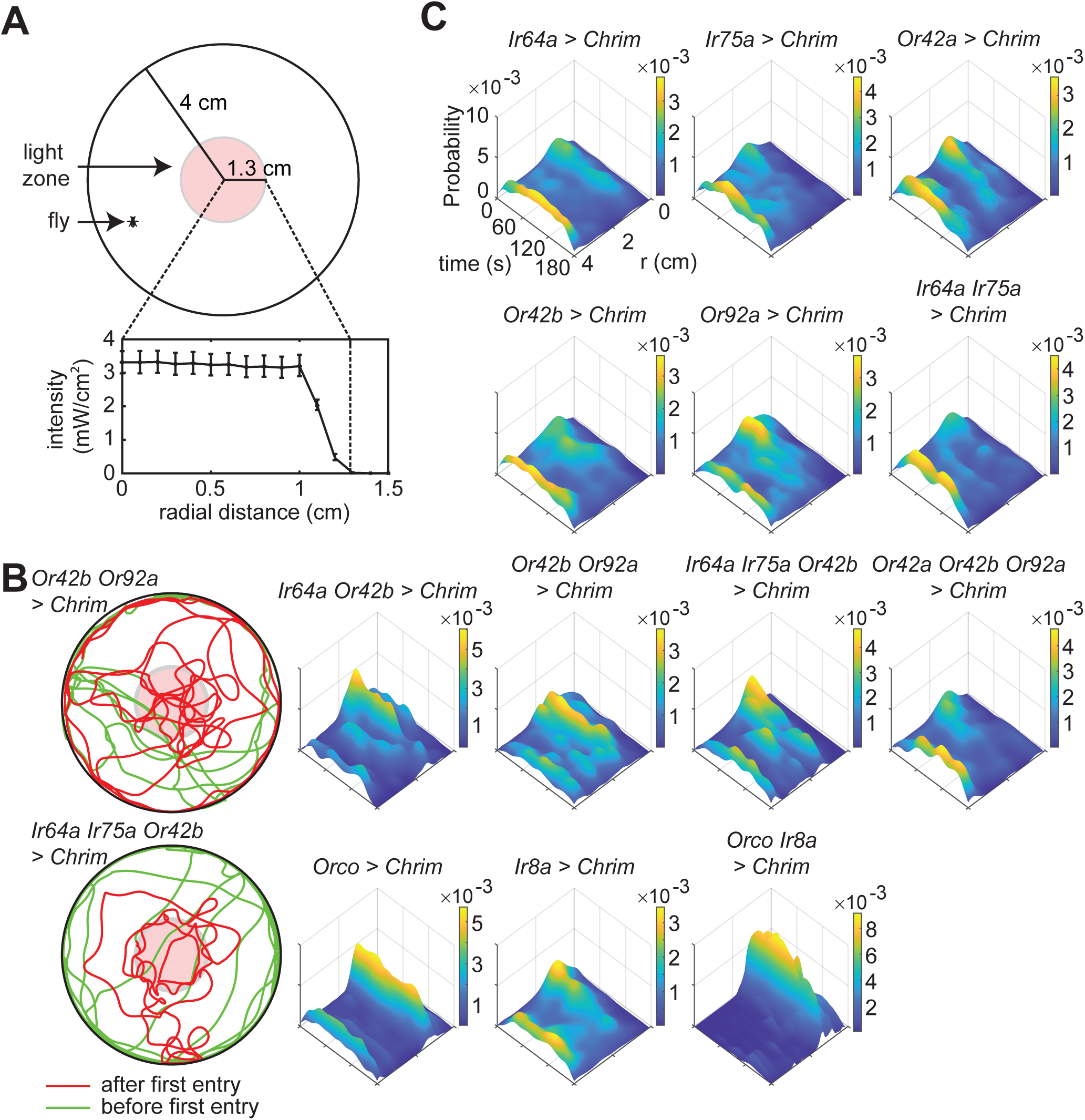
Activation of different ORN classes affect a fly’s spatial-temporal distribution in the ring arena in distinct ways. **A.** Schematic of the arena. **B.** Example trajectories showing difference between two genotypes. When *Or42b* and *Or92a* ORNs are activated, the flies spend their time more equally throughout the arena. In contrast, activation of *Ir64a*, *Ir75a* and *Or42b* results in the flies spending more time at the border. **C.** Spatio-temporal distribution of flies after first encounter with light. The plots show the probability of finding the fly at a given radial distance from the center as a function of time.

In all we activated 13 combinations of ORNs which included single ORN classes activated individually (5 different ORNs), 2 ORN classes activated in pairs (3 combinations), and combinations of three ORN classes (2 combination). (Figure 1). Because Or42b-ORNs were activated at the lowest odor concentration (Jung et al., 2015), the original experimental design was to activate two other ORNs along with Or42b – Ir64a and Or92a – to sample from ORNs that belong to different receptor classes (Silbering et al., 2011). This experimental design would imply 6 combinations in all, all of which are in this dataset. We added three more combinations to test specific ideas, none of which are relevant to our current thinking. Apart from these 9 combinations, we also activated larger sets of ORN classes by driving Chrimson under the control of *Ir8a*, *Orco*, and *Ir8a* and *Orco* together. These three receptors are co-receptors that each express in a much larger fraction of ORNs compared to olfactory receptors themselves. In all, we performed recordings from 314 retinal flies, and 289 control flies to give us 3618 minutes of data.

Activation of even a single ORN class can produce a change in the distribution of flies in the arena. Most combinations that we studied showed some change in distribution of the flies such that flies spent more time in and around the central light-zone than at the arena border (Figure 1-S1). However, each ORN combination differentially affected the fly’s spatial distribution. Two examples are shown in Figure 1B: Flies whose *Or42b* and *Or92a* neurons were activated explore the entire light-zone and make frequent forays outside the stimulated region. In contrast, flies whose *Ir64a*, *Ir75a* and *Or42b* are activated stay near the border of the light-zone. These differences in the fly’s behavior can be assessed by plotting how the fly’s density at different regions of the arena changes as a function of time after its first encounter with odor (Figure 1C). Different combinations of activated ORN classes produced different distributions at the light-zone border, within the border, or around the stimulated area. Another difference in behavior is how the behavior of the flies adapts over time (Figure 1C); behavior downstream of *Ir8a* activation adapts faster such that the flies spend less time inside at later times in the stimulus (Figure 1C, also see Figure 1-S2).

Consistent with previous work, the time spent in the vicinity of the stimulated zone when all *Orco* ORNs – consisting of 70% of all ORNs- are activated is larger than the attraction produced by a smaller number of ORNs (Bell and Wilson, 2016). Despite the large fraction of ORNs activated by *Orco*, surprisingly, when we activated both the *Orco* and *Ir8a* ORNs we observed an even larger change in behavior. Thus, it appears that the attraction produced in this arena increases with the number of ORN classes activated; this result is consistent with the idea of parallel sensorimotor transformation driving a fly’s overall behavior.

### ORN responses are shaped by locomotion making instantaneous firing rate a poor measure of sensory experience

Based on the position of the fly’s head and the intensity of light at each point in the arena (Figure 1A), we recreated the intensity of light that a fly experiences as a function of time (Figure 2A) and replayed the stimulus back to a fly in an electrophysiological rig, to measure ORN responses using single-sensillum recording, a method to perform extracellular recording from the olfactory sensilla (De Bruyne et al., 1999) that produces a slow current (referred to as local field potential or LFP) likely resulting from the opening of the channels encoded by *Chrimson*, and the resulting spiking response (Figure 2–S1). We measured responses to 6 different stimulus patterns. Based on these recordings, we created a neural encoder to predict the responses of the ORNs to any arbitrary stimulus profile (Figure 2A). The neural encoder was a cascade of two linear filters as previously described (Nagel and Wilson, 2011): the first filter described the relationship between the stimulus and LFP, and second the relationship between LFP and firing rate (Figure 2A and 2S1). There was little difference between responses from different ORN classes (Figure 2-S2). This is consistent with previous work showing that much of the difference in response dynamics result from the odor-receptor interaction, a step that the use of optogenetics bypasses (Nagel and Wilson, 2011).

**Figure 2.**
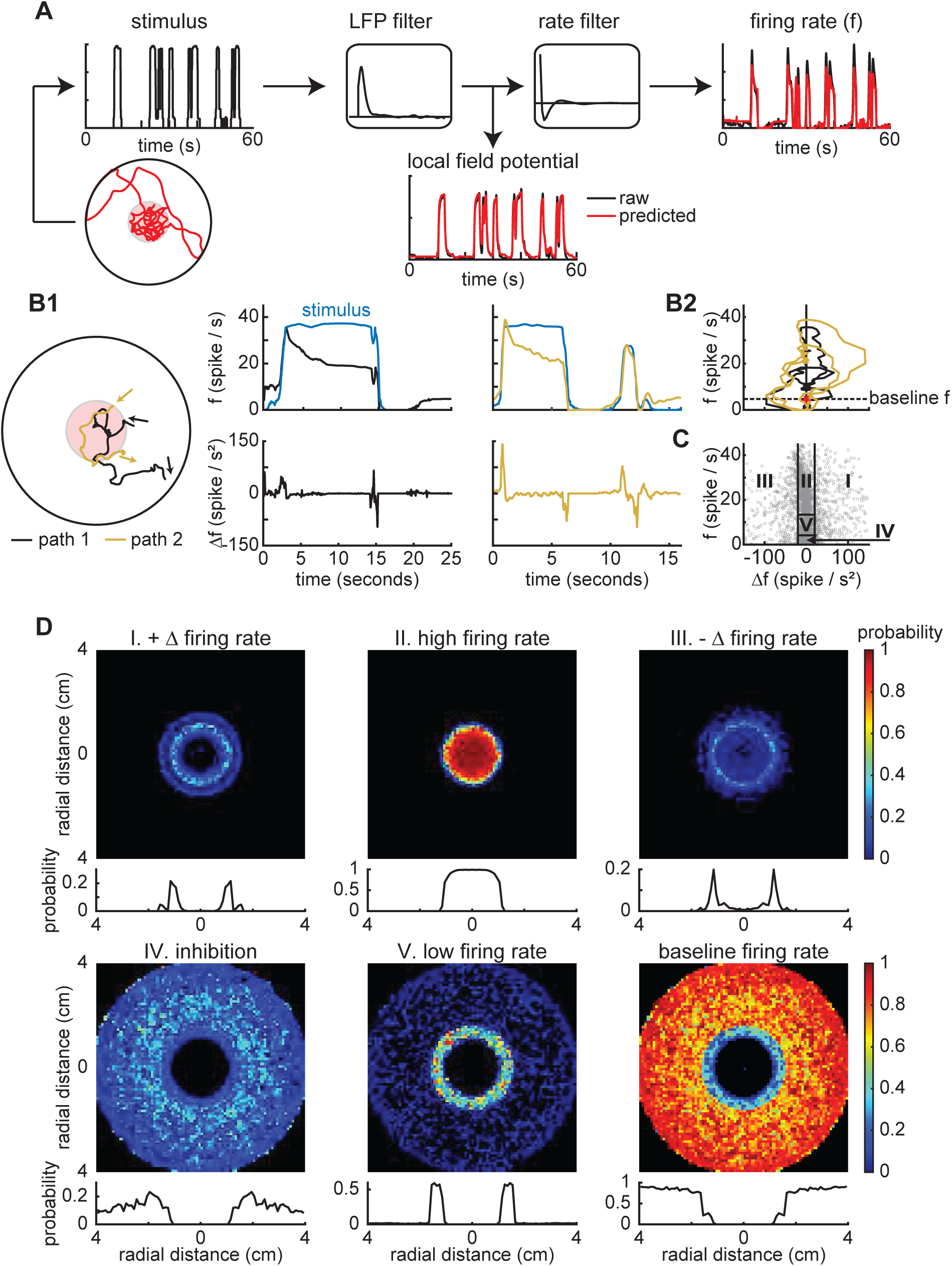
Sensory experience is approximated by instanteous ORN firing rate and change in firing rate. **A.** Linear filters were used to predict ORN responses from behavioral tracks. **B1.** Two track segments and the firing rate and change in firing rate for the black (middle) and yellow (right) show that locomotion affects ORN responses. **B2.** State space plots show the differences in sensory experience. The sensory experience is a trajectory in the *f*-Δ*f* state space. Red star indicates the beginning of each trajectory. Dotted line shows baseline firing rate. **C.** Randomly selected 10000 (gray dots) instantaneous firing rate (*f*) and change in firing rate (Δ*f*). The letters represent different regions. Although each entry into the arena results in a unique trajectory though the f-Δf space, most entries will result in sequential progression from region I to V of the state space. **D.** Top rows: The arena was split into 60x60 bins where each bin represents the probability of a state-space region (I-V) given the spatial location. Bottom rows: Probability of being in each region as a function of radial distance.

Although the stimulus is a simple step function, since the animal enters and exits the stimulus on its own, the time-course of stimulus experienced by the fly is complex. The complex temporal profile in turn makes the ORN response profile complex (Figure 2B): The ORN response when the animal transits quickly through the light-no-light boundary (Figure 2B1, yellow trace) is different from when the fly transits through this boundary slowly (Figure 2B1, black trace). Another way in which locomotion affects neural response is through their return to the arena soon after they exit. In this case, the adaptation from the last excursion to the arena affects the current response, and the peak response is lower. Differences in locomotion speed and adaptation mean that instantaneous firing rate itself is a poor measure for a fly’s sensory experience as is often assumed in studies of odor coding in immobile animal. One possible solution that an animal can employ is to measure both its instantaneous firing rate (*f*) and change in firing rate (Δ*f*). Each entry into the arena is characterized as a trajectory in the (*f* -Δ*f*) state space (Figure 2B2). Different entries into the light-zone often led to vastly different responses (Figure 2B2). These trajectories also emphasize that instantaneous firing rate is a poor indicator of the fly’s sensory experience as the ORNs have the same firing rate at multiple points along the trajectory.

Most regions in the *f* and Δ*f* space is visited, underscoring the diversity of neural response in the arena (Figure 2C). Why is instantaneous (*f*-Δ*f*) a better representation for a fly’s sensory experience? Ideally, the fly’s nervous system would encode the entire trajectory. Encoding this entire trajectory is computationally expensive, and an approximate representation of the sensory experience can be obtained from the instantaneous pair *f* and Δ*f* ; this is because although the neural response is variable, each entry into the arena goes through a similar transition starting with high firing rate and rapid increase in firing rate (region I) to high firing rate with adaptation (region II) to high firing rate with decrease in firing rate (region III) to inhibition (region IV), and finally low firing rate (V). Thus, instantaneous *f* and Δ*f* is a computationally inexpensive method for keeping track of odor history.

The usefulness of instantaneous *f* and Δ*f* is further illustrated in the spatial distribution of the flies in each of these regimes (Figure 2D), i.e., where are the flies when they are in a particular region of the *f*-Δ*f* state space? If the fly were just using firing rate, at the same radial distance near the odor border, the fly could have very different firing rate depending on whether the fly is entering in or exploring the boundary. However, taking both *f* and Δ*f* – which together describe the different regimes (I-V) - allows the flies to parse whether it is entering the stimulus region, within it, exiting the arena, or was in the arena recently.

In sum, the flies in this arena enter the stimulated zone multiple times. The instantaneous *f*-Δ*f* is a time history of its sensory experience that starts with its most recent entry to a few seconds after its exit when the ORN firing rate returns to baseline.

### Sensorimotor transformation is probabilistic, dynamic, and depends on the population of ORNs activated

Figures 1 and 2 show that instantaneous *f* and Δ*f* is a measure of the fly’s sensory experience or sensory input. The behavioral output can be described by an agent-based locomotor model (Tao et al., 2020): The agent can be in one of four states (Figure 3). Each state is described by 2-3 parameters that are chosen by the fly at each state transition and remains constant during the state. Therefore, the locomotor model is a probabilistic model with ten parameters – two of these are at the boundary so are not directly affected by ORN activation; eight parameters are affected by ORN activation (Figure 3). The effect of ORN activity is modeled as a change in distribution of these parameters (Figure 3). Specifically, *f* and Δ*f* in a short time-window before the fly transitions to a new state determines the distribution from which the fly chooses its parameter in the subsequent state. A mapping from *f-Δf* to the probability distribution of each of the eight locomotor parameters describes the sensorimotor transformation.

**Figure 3.**
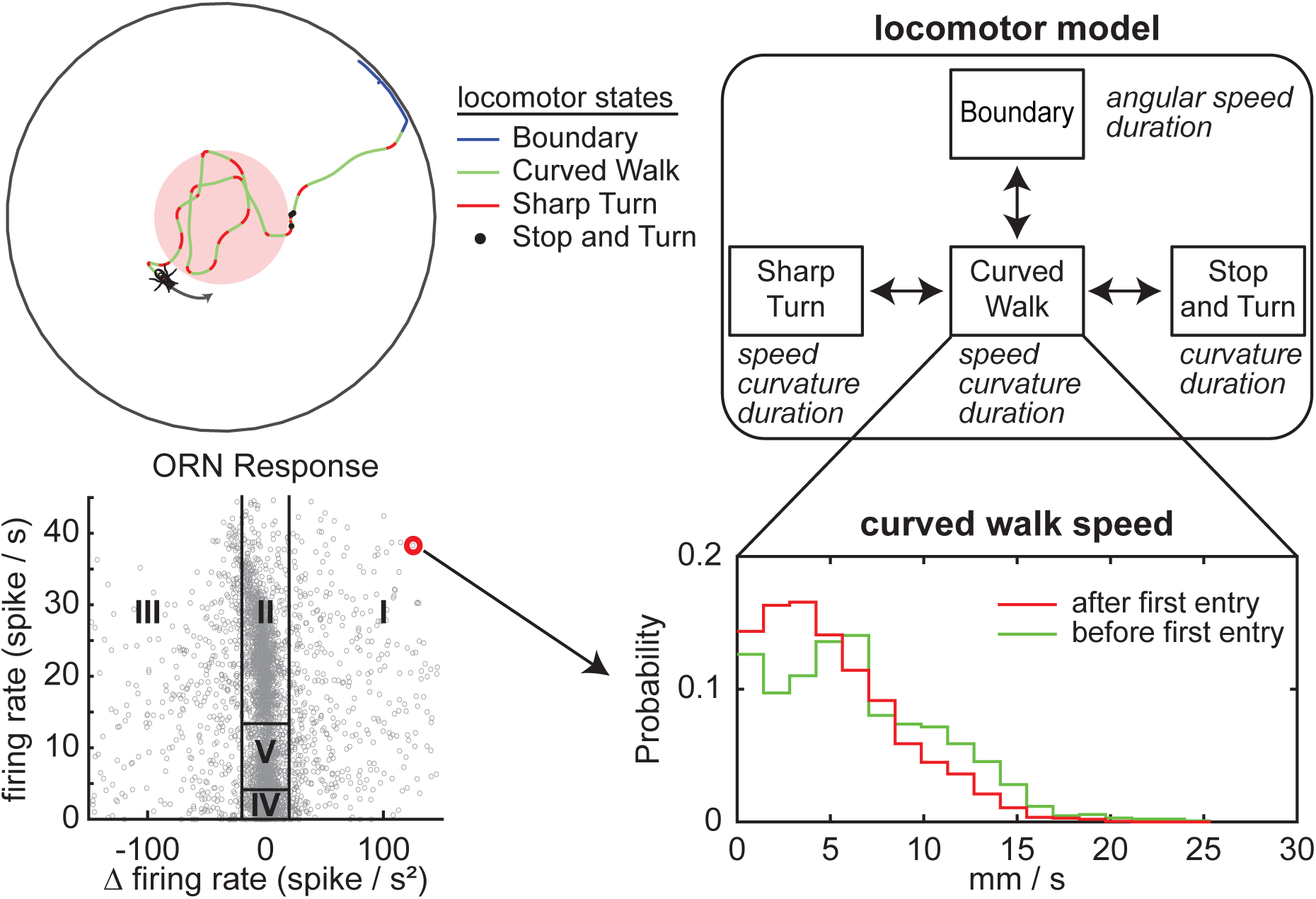
**Framework for sensorimotor transformation - instantaneous f-**Δ**f affect the distribution of locomotor parameter.** A flies’ locomotion can be described by a 4-state model of sharp turns, curved walks, stops, and the arena boundary (adapted from Tao et al., Plos Computational Biology 2020). Fly chooses the parameters for a given state at the beginning of that state from a distribution. The distribution for curved walk speed is shown. There are 8 such distributions inside the arena as well as 2 boundary distributions. Recent sensory experience - given by **f-**Δ**f** - will affect the eight distributions within the arena. Sensorimotor transformation is described as a mapping (represented by the arrow from the red point) between **f-**Δ**f** and each of these eight parameter.

To estimate how *f* and Δ*f* affects the distribution of the eight parameters, we binned all instances of the start of a state when the *f* and Δ*f* in the preceding 200 milliseconds were similar; the resulting distribution was a lognormal distribution. A lognormal distribution is defined by its mean and variance; using K-nearest neighbor (KNN) framework, we estimated mean and variance of this distribution (Figure 4-S1 and methods). In describing the effect of ORN firing on locomotion in Figure 4, we will focus on the mean of the distribution; the entire distribution is employed in generating synthetic flies (Figure 7). Despite the large number of flies that we investigated; we are still data limited in some instances. This limitation means that although we can capture the essence of the sensorimotor transformation between ORN activation and behavior, but likely miss some finer details.

**Figure 4.**
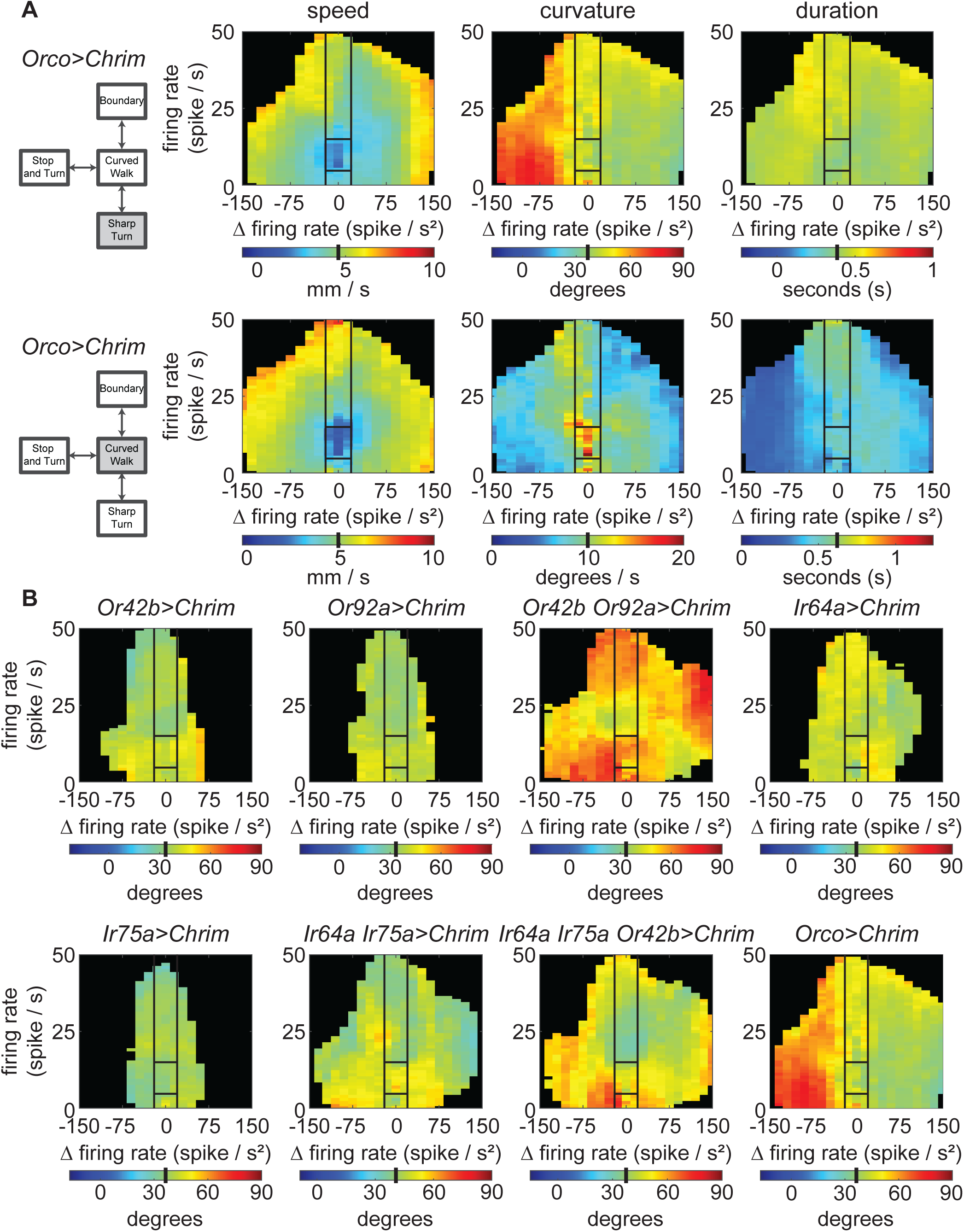
There is a unique sensorimotor mapping for each locomotor parameter. Effect of ORN activation of locomotor parameters can be described as a mapping between instantaneous f-Δf on the distribution of variables. The effect on mean is plotted here. In all panels, black lines separate out the sensorimotor mapping into 5 broad regions based on firing rate and change in firing rate. Baseline means for a given parameter before odor on is shown as a black bar in the colormap. **A.** Effect of a given combination of ORN class on locomotor parameters; sharp turn and curved walk speed, curvature, and duration. Different parameters are affected differently. **B.** Effect of different ORN classes on a given motor parameter - sharp turn curvature - is distinct.

An important feature of this sensorimotor transformation is that the effect of ORN firing depends on both *f* and Δ*f* (Figure 4A). As an example, the speed of the fly during curved walks decreases when the ORN firing rate is low (Region IV); in contrast, the speed is elevated both when the firing rate is increasing and decreasing (Regions I and III). The same effect on speed is observed during sharp turn (Figure 4A) and implies that the fly is speeding up both as it is entering and exiting the arena and slowing down while it is inside the arena and experiences a steady light intensity. In contrast to the change in speed, the largest change in sharp turn curvature occurs when the ORN firing rate is decreasing (Region III). The change in curved-walk curvature is highest in Region IV when the firing rate is steady and decreases when there are changes in firing rate. Finally, the effect on the duration of sharp turns is negligible while the curved walk durations decrease dramatically as the firing rate decreases. Other than the fact that each locomotor parameter depends on both *f* and Δ*f*, these data also suggest that the effect of a given combination of active ORNs is different on different locomotor parameters.

The effect of different ORN classes on a given motor parameter is also different (Figure 4B). As an example, consider the effect of different combination of ORN classes on sharp-turn curvature. Activation of single ORN classes *Or42b, Or92a, Ir64a or Ir75a* has a small effect on the curvature of the sharp turn. Activation of both *Or42b* and *Or92a* ORN classes together results in a large increase in the curvature of the sharp turn suggesting a strong additive effect. This additive effect is not observed in the activation of *Ir64a* and *Ir75a* together (Figure 4B). When three ORN classes - *Ir64a, Ir75a* and *Or42b* are activated together, there is a large effect on the curvature of the sharp turn suggesting that the rules of addition are and non-linear and depend on the ORN classes being co-activated. The different rules of addition are further highlighted by the effect of different ORN classes on the speed of the curved walk (Figure 4-S2). In contrast to sharp-turn curvature, activating a single ORN class – *Or42b* – has a large effect on the speed during curved walk. Activating two ORN classes has an even larger effect on speed. The effect on speed is less obvious when more ORN classes are activated (Figure 4-S2). These data are consistent with our previous work and shows that addition of more ORN classes does not simply scale the observed changes in parameter(Jung et al., 2015).

As mentioned earlier, the ORN responses following entry into the stimulus zone to a few seconds after its exit is represented as a trajectory in *f-Δf* space. A new trajectory starting at the baseline starts every time the fly enters the stimulus-zone. Thus far, we have investigated the effect of ORN firing that averages the change in behavior across multiple entries. Although the ORN firing rate returns to baseline every time the fly exits the stimulus-zone, it is possible that there are changes in the sensorimotor transformation across different entries into the stimulus-zone. To assess these dynamics, we used the KNN framework to assess how the sensorimotor transformation changes with time by evaluating the relationship between *f-Δf* and locomotor parameters at different times after the first odor encounter (Figure 4-S1B for methods). We found that the changes in locomotor parameters with time were distinct for different motor parameters; the effect of ORN activation on a given parameter can increase with time, decrease, or stay the same (Figure 4-S3). An example where the effect of ORN activation builds up over time is the speed of curved walk. The prominent decrease in the speed of walking in the arena center when both *Or42b* and *Or92a* ORNs are activated becomes more prominent with time (Figure 4-S3). In the same flies – when *Or42b* and *Or92a* ORNs are activated – there is little change in the effect of ORN activity on sharp-turn curvature over most of the duration of the stimulus (Figure 4-S3). An example where the effect of ORN activation decreases with time is the effect of *Ir8a* activation on the curvature of sharp turn; the effect of activation is largest right after the fly experiences ORN stimulation for the first time and decreases thereafter. Overall, the sensorimotor transformation for different parameters evolve differently over time.

In previous studies using both odors and optogenetics we had shown that odors or ORN activation influenced behavior not only when the stimulus is present but also outside the stimulus-zone (Jung et al., 2015; Tao et al., 2020). Some of this effect is likely due to the strong effect of the ORN off-response (Region IV). Is the effect of ORN activation persistent after the ORN firing rate has returned to baseline?

Indeed, when we analyzed kinematics only during the period when the firing rate was at baseline levels, we found that for some genotypes there was no change in kinematics during the baseline period (Figure 4- S4). For other genotypes there are changes in kinematics even when the ORN firing rate has returned to baseline (Figure 4-S4). These changes also adapt over time.

### Flies can turn preferentially at the border to either stay inside or turn inwards

Apart from kinematics, insects can use directional information to direct their turns so that they turn towards the stimulated region. We have already shown that flies use directional information in the ring arena (Tao et al., 2020). This use of directional information is clear in the tracks of flies (Figure 5A) which show flies weaving in and out of the light-zone suggesting that they use directional information. We quantify the extent to which flies use directional information by quantifying turn optimality (see methods). As the fly exits the light-zone, unless it is moving exactly in a direction normal to the light-zone, it can either turn in a direction where a small turn will bring it inwards (optimal direction) or in a direction in which a large turn is necessary (Figure 5B). Similarly, as the flies enter inside, the flies can turn in an optimal or non-optimal direction (Figure 5B). Indeed, flies show a preference for turns in an optimal direction (Figure 5C). They turn towards the border as they exit the arena with a higher than chance probability. In contrast, the flies turn back into the light-zone when they are inside.

**Figure 5.**
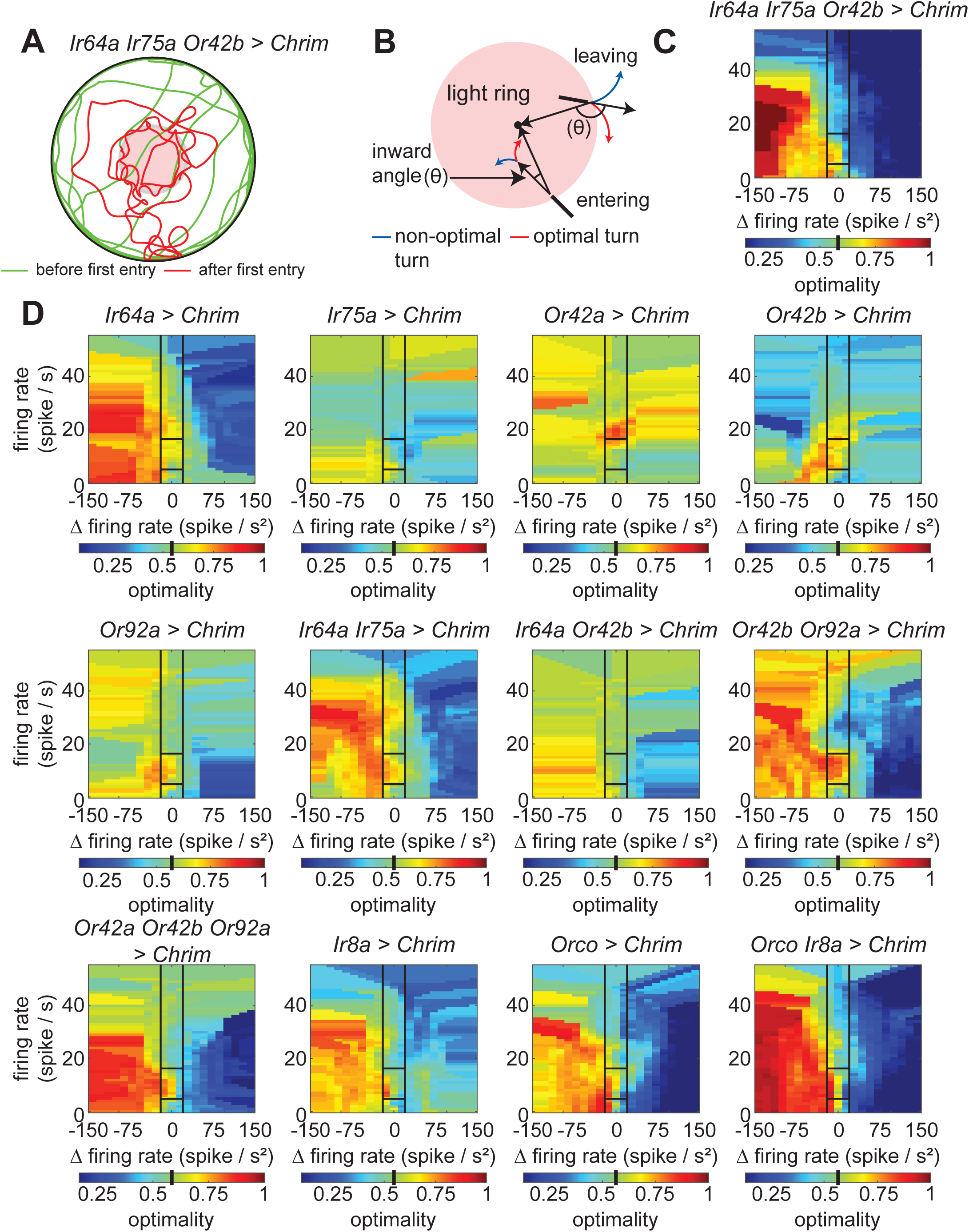
ORN activation alone allows flies to perform directed odor tracking. **A.** A sample behavioral trajectory shows that flies are able to track the stimulus border. **B.** Schematic showing optimal and non-optimal turns. Turn direction that decreases the angle between the current direction of the fly and radial vector (or θ) are optimal. **C.** Turn optimality (defined as the percentage of optimal turns) for *Ir64a Ir75a Or42b* > *Chrimson* relative to baseline. Black lines separate out the sensorimotor mapping into 5 broad regions based on firing rate and change in firing rate. Baseline turn optimality is shown as a black bar in the colorbar. **D.** Turn optimality for all other combinations of active ORN classes showing that different ORN classes affect optimality in different ways.

This directional preference also depends on genotype. When a single ORN class is activated, the flies do not exhibit this directional preference except when Ir64a-ORNs are activated (Figure 5D). Consistent with complex rules of integration observed for kinematic parameters, when both Ir64a-ORNs and Or42b- ORNs are activated, the directional preference becomes smaller. All other combinations in which two ORN classes are activated show an increased propensity for directed turns. Finally, consistent with the border-hugging tracks when *Ir64a, Ir75b and Or92a* ORNs are activated, turn bias is particularly strong when this genotype is activated (Figure 5C).

### A model for diverse rules of integration between ORN activation and change in locomotor parameters

It is clear from the data presented in Figures 4 and 5 that ORN activation affect multiple motor parameters in a manner that is dependent on both *f* and Δ*f* and on the identity of the active ORNs. Given that there are both excitatory and inhibitory PNs that are each uniglomerular and multiglomerular; and there is considerable integration between PN-types within the lateral horn, the neural substrate for parallel sensorimotor transformation clearly exists (Schlegel et al., 2021). In this section we develop a conceptual framework for this parallel sensorimotor integration. We propose that signals from each ORN is carried through parallel channels that either invert the signal, send them along without inversion; are either differentiating or integrating, together form five output channels downstream of each ORNs that correspond to the five regions of the *f-Δf* space (Figure 6-S1). Each entry into the stimulus-zone will lead to a sequential activation of these output channels. Each channel can have a different effect on a given motor parameter that can be modeled as a change in the distribution (Figure 6-S1). Signals from different ORN classes are integrated according to different rules in each of these five output regions. The rules of integration were modeled using a regression model (Figure 6-S2).

**Figure 6.**
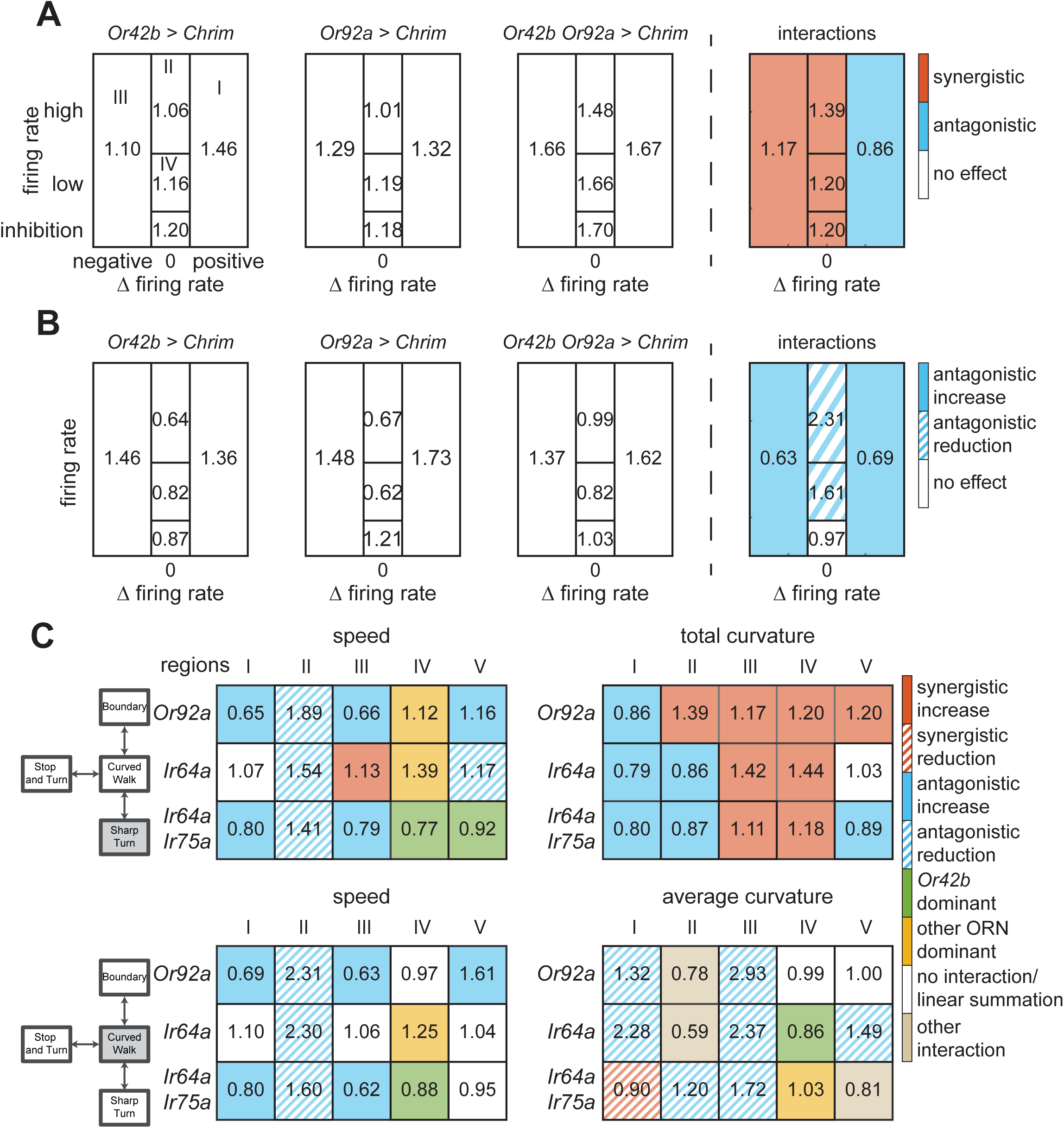
Rules of integration between ORN classes are diverse and depend on region of the state-space and the locomotor parameter. **A.** Rules of integration for the effect of *Or42b*, *Or92a* on sharp turn total curvature. The three panel on the left are the effect on sharp turn. 1= no effect; 1.46 is a 46% increase. The rightmost panel shows the interaction. Color implies that the interaction was synergistic such that the observed effect is 10% larger than for linear summation (orange), and 10% smaller (blue). **B.** Same as **A**, but for curved walk speed. The same two ORN class can have either synergistic or antagonistic interaction depending on the locomotor parameter. Synergistic and antagonistic interactions can be further delineated by whether the individual ORN activation results in an increase (solid) or decrease (hashed) in locomotor parameter. **C.** Interactions between *Or42b* and three other ORN combination *Or92a*, *Ir64a*, and *Ir64a;Ir75a* show that rules of summation between the same ORN classes are diverse. Numbers represent interaction effects on locomotion.

We applied this approach to all combinations of ORN classes and illustrate this analysis with how activities from *Or42b-ORNs* and *Or92a-ORNs* affect sharp turn curvature (Figure 6A). Activation of *Or42b-ORNs* has a large effect (>40% increase in Region I) as the fly enters the arena. This effect is transient as reflected by the small effect in Region II when the fly is fully inside. The effect on curvature returns when the fly is exploring the odor border (Region IV), and when the ORN response is inhibited (V). Activation of *Or92a-ORN* alone also has a similar effect. When the two ORNs are activated together, there is a large synergistic effect on the curvature of sharp turns except in region I. The effect of this synergism is that the sharp turn curvature is high in all regions when both ORNs are activated. The synergy between the two ORNs mean that the effect due to the two ORN classes together is 20-40% higher than what would be expected from a linear sum (Figure 6A, rightmost panel). Interestingly, although there is still a large effect of the combined ORN activation on sharp turns when the fly is entering the arena (Region R1), the interaction effect is not synergistic as the observed effect of the two ORN classes is about 15% smaller than what would be expected from a linear summation.

Interaction effects can be diverse even for the same two ORNs effect on a motor parameter: Consider the effect of *Or42b* and *Or92a* ORNs on curved walk speed. In some regions (I, III, IV), the effect of the two ORNs add sub-linearly such that the increase in speed is not as large as expected from a linear summation. The large decrease in speed observed in region II when individual ORN classes are activated is completely abolished when both ORN are activated (Figure 6B). Interaction terms can also be such that the effect of one ORN class dominates over the other particularly when the two ORN classes individually have different effects, or can affect individual parameters when neither has an effect individually (not in this case, see Figure 6C for an example).

This diversity of rules is evident in the effect of *Or42b-ORNs* when combined with different ORN classes (Figure 6C). For a given parameter, the effect depends on the region of the *f-Δf* space. As an example, take either the curved walk or sharp turn speed: In region II, the effect of activating *Or42b* in combination with any of the other ORNs is to increase the speed. This is even though *Or42b* activation by itself reduces the speed in region II. In contrast, in regions I and III, Or42b activation results in a decrease in speed. Finally, during the inhibition epoch (region V), the effect of *Or42b* appears muted and the overall behavior is close to the behavior upon activation of *Ir64a-ORNs*.

The same diversity applies to the effect on curvature and the effect is different both for different regions of *f-Δf* space and for sharp turn and curved walk curvature: In regions I and III, i.e., when the flies are leaving or entering, *Or42b* activation has an antagonistic effect on the curved walk curvature such that the summed effect is smaller than what would be effected from a linear sum (Figure 6C, bottom right panel, hashed blue). In many cases, the net effect of activating Or42b is so strong that the change in curvature due to Or92a or Ir64a activated alone is almost abolished. In contrast, the sharp turn curvature increases in many regions when Or42b is activated along with the other combinations.

This diversity of integration rules and its dependence on both f-Δf makes sense in the light of the fact that the motor program should change with the fly’s sensory experience. This diversity can not only be supported by olfactory processing circuits but is the only possibility given the widespread convergence and divergence of olfactory signals in higher olfactory circuits. If we follow only the most salient, feedforward connections, Or42b-ORNs signal to at least 5 different LHONs which all integrate input from different populations of ORN classes (Figure 6-S3).

### A generative model for the effect of odors on locomotion

The analysis above (Figures 4-6) shows that the effect of ORN activation on locomotion depends on the identity of the ORNs activated. Can the changes in kinematics explain the changes in distribution of the flies? To this end, we created synthetic flies based on our agent-based model(Tao et al., 2020). The details are in the Materials and Methods section and in Figure 7-S1. Briefly, just like the experimental flies, each synthetic fly walked for 6 minutes - 3 minutes before the light turned on, and three minutes following light on. Synthetic flies start at the center of the arena and moved around the arena through a series of transitions into the four states. Synthetic flies always started in curved walk. Curved walk ended in stop, sharp turn or at the boundary (Figure 3A). Tracks corresponding to each transition was generated by sampling from speed, curvature, and duration distributions for each state for the *f* and Δ*f* during the 200 milliseconds preceding each state transition. Essentially, using the position of the synthetic flies as a function of time, we estimate the intensity of light experienced by the fly as a function of time; this light intensity was converted into ORN spike rate using the two-stage linear encoder derived in Figure 2. The resulting *f* and Δ*f* was used to determine the distributions that the synthetic flies sampled from at any given time. The duration that each transition lasted was also selected from the empirical distribution (Figure 4).

To assess how well the behavior of the synthetic flies replicated that of the empirical flies, we first compared the flies whose *Orco-ORNs* are activated, a genotype that we have analyzed previously (Tao et al., 2020), and because these flies show a large change in their distribution. The radial distribution of the empirical and synthetic flies before and after the central red light is turned on is shown in Figure 7A. As expected, the radial distribution of the synthetic and empirical flies before the light is turned on is similar. The distributions of the empirical and synthetic flies during the stimulation period are clearly different (Figure 7A). This mismatch is not due to large differences in kinematics as much of the kinematic changes in the empirical flies are replicated in the synthetic flies (Figure 7-S2).

**Figure 7.**
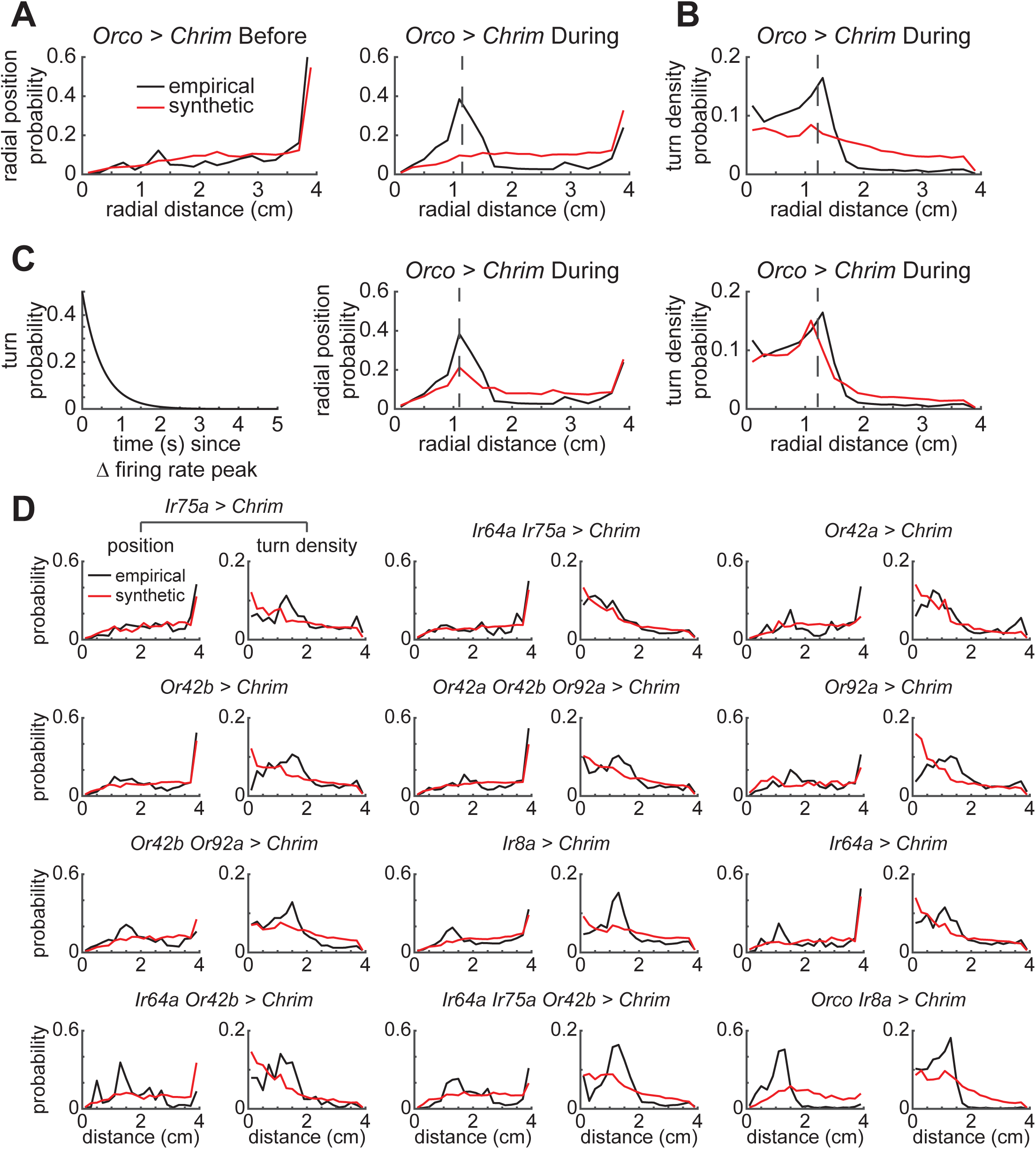
An agent based model of flies based on locomotor kinematics and turn optimality can capture much, but not all of the flies behavior. A. Spatial distribution of the empirical and synthetic flies are similar before the stimulus is turned on. During stimulation, the synthetic flies spend less time than the empirical flies inside the stimulated area but more time between the stimulated zone (dashed line) and the outer boundary. B. The difference in radial distribution results from a larger propensity to turn at the border of the stimulated zone than the synthetic flies. C. An exponentially decaying probability of transitioning into a turn was used to model the empirical turn density (right). By improving the turn density fit, the model was capable of better capturing the spatial position (middle) of flies in the arena. D. When there is no or small increase in turn density at the border, the synthetic flies have a distribution close to the empirical flies. Examples are *Ir64a* alone, and *Ir64a* and *Ir75a* activated together. As the turn density increases, the pattern observed in the *Orco* activated flies is observed.

A closer examination of the radial density in Figure 7A shows that the crucial difference is that the synthetic flies spend more time in the region between the outer border and the light-zone than do the empirical flies and lack the large peak in radial density at the border of the stimulated region. These differences imply that the kinematics we measured do not fully capture the behavior of the flies at the border of the light-zone. This result is consistent with our previous work where an assumption that every time the fly is at the border, there is a change in state was necessary to fit the data(Tao et al., 2020).

Indeed, we found that a negative peak in Δ*f* caused a large increase in the propensity of the flies to exit curved walk and enter sharp turn (Figure 7-S3). Although our measurement of kinematics does capture this tendency - the curved walk duration is lower when Δ*f* is negative (Figure 4A) - this tendency is underestimated because of estimation errors in the KNN model originating in the lack of data for high negative Δ*f*. Consistent with this idea, there is a large peak in the propensity to turn in the empirical flies that is not captured in the synthetic flies (Figure 7B). We attempted to recreate this increased turn propensity by adding a “border choice” parameter. Although, we were not completely successful in matching the turn density observed in empirical flies, we show that incorporating the increased turn density in the model for synthetic flies results in a large peak in the radial density near the light-zone and decreases the density in the unstimulated, non-border regions (Figure 7C).

The trend observed in *Orco-ORNs* activated flies is observed in other genotypes. In flies in which there is small peak in turn density show a close match between radial density in the empirical and synthetic flies (Figure 7D). Combinations of ORNs that result in increased turn density show the same trend as seen in the Orco-ORNs – peak radial density at the border of light-zone that is not captured in the synthetic flies and increased density in the synthetic flies just outside the light-zone. A formal analysis confirms the trend (Figure 7-S4) and shows that the difference between the radial density for empirical and synthetic flies is correlated to the increased turn density at the border of the stimulated region.

## Discussion

### A probabilistic model describes the sensorimotor transformation in the olfactory system

The behavior of a fly walking around in a small arena can be approximated as a series of discrete choices: the fly chooses a speed and a curvature and walks at the same speed and curvature for a given duration before making another choice. Based on its choices, our locomotor model has four states: curved walk, sharp turns, stops, and boundary walking. The selection of locomotor parameter is probabilistic: it chooses a speed, curvature, and duration (or locomotor variables) from a distribution. Thus, the fly walking around an arena sequentially transitions from one state to another choosing its motor parameters from a distribution. ORN activation has two effects on behavior. First, ORN activation changes the distribution from which the fly selects its speed, curvature, and duration. Second, because ORNs are activated only in the stimulated region, this creates an asymmetry; flies take advantage of this asymmetry by biasing their turns such that each turn directs them towards the stimulus zone.

The fly is in a dark arena without any strong visual cues. Stimulating the fly’s olfactory system also does not provide strong directional cues; locomotion makes sensory assessment even more uncertain as sensory delays mean that the current firing rate is only reporting events that occurred at a different location and in the past. Under these conditions where there are few navigational cues, a probabilistic approach to locomotion is appropriate. While the exact nature of the locomotor model and how sensory information including olfactory information affects the model parameters depend on the species, the size of the arena, the nature of the stimulated area, we expect that a simple probabilistic model will always capture the essence of the underlying sensorimotor transformation because most real-world locomotion problems are solved using simple strategies under conditions where animals are information-starved (Wang and Hayden, 2021).

### Dependence of locomotor parameters on f-Δf pair is a mechanism for automatically adapting locomotion to sensory context

A common approach to sensorimotor transformation is to create a filter that relates firing rate to some behavioral parameter. A recent study in the context of olfactory behavior also uses the same basic conceptual framework (Álvarez-Salvado et al., 2018). In most of these studies, behavior is performed in an open loop, i.e., although sensory stimulus can shape motor output, the reverse is not true. In nature, most behaviors are performed in closed loop where sensory stimulus affects behavior which in turn affects the stimulus experienced at the next moment. A consequence of locomotion controlling sensory experience is that the same stimulus will yield a different response depending on the stimulus history which in turn depends on locomotion. Therefore, using firing rate to control locomotor parameter is not an appropriate strategy, and indeed it is clear that the ORN firing rate is not transformed into changes in locomotor parameter. At the same ORN firing rate, different rate of change in firing rate had different and sometimes opposite effect on a locomotor parameter (Figure 4).

Ideally, neural circuits would integrate information across time to create an internal representation of sensory experience. Some integration across time is indeed possible but most models of olfactory search are based on modulating behavior based on instantaneous sensory experience (Vergassola et al., 2007). It is also unclear how information sources necessary for guiding locomotion– such as sensory information, location, and speed would be integrated together. We show here that instantaneous *f* and Δ*f* can serve as an adequate surrogate for integrating information across time: Together they can signal whether the fly is going up or down concentration gradient or is in an area with steady stimulus concentration without any need to integrate information across time. In other words, instantaneous *f* and Δ*f* can together serve as short-term memory from the time the fly enters the stimulated zone and until it leaves.

Using instantaneous *f* and Δ*f* to modify behavioral parameters is an effective and simple strategy to continuously adapt motor program to sensory context. Take for example, the change in speed. As the fly enters inside the stimulus-zone and its ORNs are increasingly more active, the fly starts moving faster. Conversely, as it enters a region of constant stimulus and the changes in firing rate is low, the speed decreases. The relation between speed and firing rate change might reflect a change in locomotion strategy; increased speed as the fly is moving up the concentration gradient changes to a search behavior with low-speed careful search of an area when the firing rate is constant. Basing behavior on *f* and Δ*f* is a mechanism for automatically modulating it based on previous sensory experience.

The instantaneous *f* and Δ*f* can only signal short-term context – from the current entry into the stimulated region to exit. On a longer timescale, the sensorimotor mapping between *f* and Δ*f* and locomotor parameters is altered and allows modulation of locomotion over successive encounters.

### Different relationship between f-Δf pair and locomotor parameters is another mechanism for automatically adapting to sensory context

In all, there are eight kinematic parameters; out of these 8 parameters, all but the duration of sharp turns is affected by ORN activation. Out of the remaining seven, the speed of curved walk and sharp turns are affected similarly across all genotypes and is likely a single parameter, leaving six parameters that all show different dependence on *f* and Δ*f* and on combination of ORN class activated. In this section, we discuss that the distinct relationship between *f* and Δ*f* locomotor parameter is another facet of sensorimotor mapping that ensures a motor program that automatically adapts to sensory context.

The observed change in speed – increases when there are large changes in ORN firing rate, and a decrease when the firing rate is elevated but constant is consistent with our previous work(Jung et al., 2015). The decrease in speed is a hallmark of search behavior observed after an animal finds a resource or an odor that indicates the resource. Decrease in curved walk speed is often – but not always – associated with an increase in curvature; this decreased speed and increased curvature is a simple, conserved strategy that allows an animal to stay close to the stimulated region; a phenomenon that is also observed in field studies (Bergerot et al., 2013). Activation of single ORN classes – *Or42b*, *Or42a*, *Or92a*, *Ir64a*, and *Or42a* and to a lesser extent Ir75a – all result in this pattern of speed change; activation of combinations also produces the same change. These results are consistent with the idea that activation of most ORN classes would result in a change in curved walk speed and curvature that initiates a local search.

Contrasting the dependence of curved walk speed and curvature versus sharp turn curvature illustrates the logic of the underlying sensorimotor transformation. Sharp turn curvature changes most as the firing rate decreases while curved walk speed/curvature changes when the changes in firing rates are small. These changes are consistent with the idea that increase in sharp turn curvature are employed when the fly is exiting the stimulated region and wants to return. Once again, a mapping between instantaneous *f* and *Δf* and sharp turn curvature accomplishes a part of the overall motor program without the necessity of temporal comparisons. An increase in sharp turn curvature is a conserved strategy documented in other insects (Sabelis et al., 1984; Waage, 1978) which also increase their turn amplitude to return to a resource patch. In a previous study we have shown that this increase in turn amplitude is important for a fly spending more time in the stimulated region(Tao et al., 2020). Although single ORN classes can cause some change in sharp turn curvature, the rules of integration across ORN classes are quite different from the integration rules for curved walk speed and curvature.

The difference between dependence of the duration of curved walk on combination of ORN and that of sharp turn further emphasizes the idea that these motor programs are modulated by parallel sensorimotor circuits. As the fly exits the stimulated region, there is a large decrease in the duration of curved walk.

This change in curved walk duration means that as soon as the firing rate starts to decline, the flies stop walking and make a turn. It would make sense if this decrease in curved walk duration was strongly coupled to the increase in sharp turn curvature. Surprisingly, changes in curved walk duration requires the activation of many more ORN classes than the increase in sharp turn curvature implying that they are likely to be modulated by parallel sensory circuits.

The distinct dependence of different locomotor parameters on *f* and *Δf* is easy to conceptualize as a description of a motor strategy that unfolds as the sensory context changes. The distinct dependence on different ORN classes is more difficult to rationalize. It is possible that there is no clear rationale, and that this dependence is a reflection of how these circuits evolved over millions of years building different sensorimotor loops. Having access to multiple sensorimotor loops ensures that a fly will find its resource in any environment.

### Implications for circuit mechanisms for sensorimotor integration in the olfactory system

At first glance, the behavioral strategy we outline above seems complicated and raises the question whether neural circuits in the fly’s brain can execute this strategy. A closer look at the olfactory circuit in the fly brain suggests otherwise; the circuit architecture underlying the signal propagation from the ORNs to the antennal lobe and further to the lateral horn is well-suited to perform this computation. To execute the behavioral strategy outlined above, these circuits must perform two computations. First, the circuits should be able to compute instantaneous f-Δf. Computing f-Δf means that different higher-order neurons respond most strongly to different phases (such as rising phase, plateau etc.). This differential response is possible because each ORN class connects to multiple second-order neurons called the projection neurons (PNs). There are ∼350 PNs in all, ∼200 PNs are cholinergic and excitatory; ∼100 PNs are GABAergic and inhibitory; neurotransmitters for the rest are uncertain (Bates et al., 2020). The PNs are divided almost equally between uniglomerular PNs that send their dendrites to a single glomerulus, and multiglomerular PNs that project to multiple glomeruli. The presence of almost equal numbers of excitatory and inhibitory PNs imply that it would be easy to perform sign inversion in higher-order circuits. Differences in kinetics is also likely. The most well-studied PN class – excitatory uniglomerular PNs – act as differentiators owing to the synaptic depression at the ORN-PN synapse (Kazama and Wilson, 2008). It is possible that other classes of PNs in the antennal lobe has different kinetics as the inputs into different PNs within each glomerulus is heterogenous. Functional heterogeneity in PN responses within a glomeruli has been observed in other insects (Baker and Hansson, 2016; Lee et al., 2019).

Second, our data suggests that activities from two ORN classes are integrated according to different rules based on both instantaneous f-Δf and the locomotor parameters. This integration can happen both at the level of mPNs and at the level of lateral horn. Each glomerulus makes connection with ∼30-40 mPNs; the number and diversity of mPNs imply that there is enough neural circuitry to compute different rules of integration. Similarly, there is great diversity in cell-types in the lateral horn. Based on this connectivity pattern, the lateral horn consists of ∼500 cell-types in Drosophila (Schlegel et al., 2021). There are also >37 types of output neurons from the lateral horn.

### Implications for attraction versus repulsion

As already mentioned, much of the work in flies is aimed at understanding whether a given ORN class is attractive or repulsive (Semmelhack and Wang, 2009; Tumkaya et al., 2022) or the valence of a given ORN class. What are the implications of parallel sensorimotor loops on the valence of a ORN class?

Since single ORN classes almost always cause a change in speed, we expect that activation of single ORN class could result in some attraction, particularly in arenas where they can return to the stimulated region frequently. However, attraction is not guaranteed even when there is a change in speed. Even in our current study, if we define attraction as fraction time spent in the stimulated region, attraction downstream of single ORN classes is small (Figure 1-S1) as the fly spends more time in the vicinity of the stimulated region rather than within the region itself. This result – that activation of single ORN class would result in either no attraction or mild attraction – is consistent with most work on attraction (Tumkaya et al., 2022).

As more ORN classes are activated, more of the motor repertoire is recruited leading to greater attraction. One exception is that when *Or42b,Or42a* and *Or92a* are all activated, the flies are less attracted than when *Or42b* and *Or92a* are activated. Does this mean that Or42a is a repulsive ORN class? This is not the case as activation of *Or42a* by itself is mildly attractive. If we think of the propensity of a fly to take the optimal turn as a measure of its intention to return to the odor, then *Or42b,Or42a* and *Or92a* active flies should be very attractive as they shows strong turn bias (Figure 5). It appears that this combination of activated ORN classes cause smaller changes in curved walk speed and duration, and smaller change in sharp total curvature likely leading to less time spent near the stimulated region. One interpretation is that an intent to be attracted to the stimulated region does not necessarily mean that the sensorimotor transformation necessary to achieve that intent is a given.

A surprising finding from multiple studies is that activation of all *Orco*-ORNs (Larsson et al., 2004a) – which consists of 70% of all ORNs – results in a large attraction. Here we add to this surprising finding by showing that activation of *Orco* and *Ir8a* ORNs together results in an even stronger attraction. These data are inconsistent with the idea that some ORNs mediate attraction and others repulsion, or with the idea that ORN population activity signal particular objects that are either attractive or repulsive. There are two possible interpretations. First, these data are consistent with the idea that different ORNs modulate different locomotor parameter, and more ORNs that are active, more efficiently these motor parameters are activated resulting in greater attraction. At the same time, it is important to note that activating a combination of three ORN classes – *Ir64a*, *Ir75a*, and *Or42b* – mediates robust attraction. Second, it is possible that there are some repulsive ORN classes that elicit motor programs that would result in the flies moving away from the stimulated zone, but their effect is overwhelmed by ORNs that mediate motor programs that lead to attraction.

### Implications for navigation and working memory

There are two changes in a fly’s locomotion. First, are the instantaneous kinematic changes that we presented in Figure 4 and discussed in the previous discussion section. Second are memory-driven changes – these are changes that require a fly to accumulate evidence over time or remember past sensory experience. We observe three such changes: First, there are some changes in behavior even after the ORN firing rate reaches baseline. Among these changes are the fact that the arena border is less attractive, i.e., the stimulated fly spends less time at the arena border. There are kinematic changes as well even when the ORN firing rate is at a baseline (Figure 4-S4). These effects represent short-term memory that lasts approximately ten seconds. These short-term memory effects are likely mediated by dopamine-mediated circuit modifications in the fly’s mushroom body which are capable of modifying behavior on a short time-scale (Cohn et al., 2015; Modi et al., 2020).

Second, the instantaneous kinematic changes themselves are subject to change with time after the first exposure to the stimulus (Figure 4-S3). Although we do not have enough data to completely characterize the dynamics of these changes, these dynamics depend on the ORN class being studied and the locomotor parameters being studied. One prominent adaptation is that as time since first stimulus encounter increases, flies spend more time at lower speeds and higher curvature inside the stimulated area consistent with a more intense search. These behavioral changes are also likely to be mediated by dopamine mediated circuit modification in the mushroom body and signaling to both lateral horn and downstream motor circuits by mushroom body output neurons (Frechter et al., 2019; Schlegel et al., 2021).

Finally, flies turn in the optimal direction as they exit the arena. This turning in the optimal direction is elicited by a large drop in the rate of ORN firing but it likely implies that the fly has some spatial sense of the stimulated area and its own locomotion with respect to this area. This behavior is reminiscent of similar behavior reported in the presence of a drop of sugar (Kim and Dickinson, 2017); the only difference is that the behavior in our arena is triggered by a stimulus instead of purely through navigational cues.

### Implications for control of continuous real-world behavior

Much of the work on the neural basis of behavior has been performed on discrete behavior where the animal is making a binary choice or a choice between a small number of options. Discrete behaviors are self-contained – the choice is irrevocable and does not affect future choices; often the animal has a relatively long time to make a decision. In this framework, nervous system is an information processing organ (Marr, 2010; Riesenhuber and Poggio, 1999): it constructs increasingly sophisticated and abstract internal representations of the world. These representations are stored in working memory to make decisions, manipulate representations to build complex knowledge, and to recruit the motor system to perform behavior. This serial model for behavior works well for discrete behaviors. This framework has its origins in psychology, as well as in work performed in the context of complex human abilities of abstract problem solving (Newell and Simon, 1972). Many of the decisions that we make in our life and perhaps dominate so much of our conscious mental life are also discrete. Because it makes sense, and because it is still the dominant model, a student of neuroscience would be strongly inclined to this model after picking up any neuroscience textbook (Kandel et al., 2000; Purves, 2008).

However, many of our behaviors are not discrete. Walking to a car, making a peanut butter sandwich are all continuous behavior which requires continuous sensorimotor integration. These actions require constant interactions with a complex environment. At each instant, there is a bewildering array of choices instead of a single choice. Moreover, each choice does not result in a final outcome. Although a study of real-time, natural behavior has been the foundation of ethological research for a long time (Hinde, 1966), because of technical challenges their underlying neural mechanisms are seldom studied. In the absence of studies aimed at understanding the neural underpinnings of such continuous behaviors, many studies have assumed that the serial processing approach that works for discrete behavior also applies to continuous behaviors. This assumption has been challenged on multiple fronts: There is no region in the brain that integrates all the information (Cisek and Kalaska, 2010; Colby and Goldberg, 1999); most brain regions are strongly affected by on-going motor activity (Allen et al., 2019; Gründemann et al., 2019; Stringer et al., 2019); perception, action-selection, and execution of behavior occur in parallel rather than serially (Cisek and Kalaska, 2010); circuits in the brain are reciprocally interconnected making serial processing unlikely (Hegde and Felleman, 2007; Zeki, 2018); efforts to make robots that perform continuous behavior based on the serial model have not been very successful (Brooks, 1991).

These considerations have led researchers to further develop alternative models for continuous behaviors; models that have already existed in neuroethology. All these models propose a modular organization containing parallel sensorimotor loops. In many cases, each of these loops represents a solution to one aspect of an ecological problem (Wehner, 1987; Wessnitzer and Webb, 2006). In these models there is no strict temporal hierarchy between action selection and its execution. Instead, the two occur in parallel; the environment dictates the palate of actions at any moment. Action selection occurs gradually as action execution slowly reduces the palate to a single action (Cisek, 2007). In these models, an internal representation of the world is not necessary. The tight coupling between the processes between sensation and action makes assigning causal relationship between one process and the next impossible.

Similarly, modularity – the idea that different circuits in the brain perform different functions - is also not valid in these models.

Our data supports a modular organization with parallel sensory motor loops and provides a granular model for continuous behavior. Our results are best interpreted in a control theory framework (Gallivan et al., 2018) in the context of multi-step behavior. The fly is endowed with a set of controls – the parameters of the locomotor model. These parameters are controlled by the state of the system – defined by *f* and Δ*f* – through a control policy. The mapping between *f* and Δ*f* is the control policy. The goal of the control is to stay close to the stimulated region and search the stimulated region thoroughly. On a longer timescale, this mapping is constantly updated as the relationship between *f* and Δ*f* and different locomotor parameters is plastic; this plasticity allows both the goal and control policy to adapt as necessary.

## Methods

### Flies

Flies were raised in sparse culture conditions consisting of 50 mL bottles of standard cornmeal media with 100-150 progeny/bottle (Bhandawat et al., 2010). Active dry yeast was sprinkled on each bottle after removing the parents (1-3 days) to enrich the larvae’s diet. Bottles were placed in incubators set at 25°C on a 12hr dark/ 12hr light cycle. 10-15 newly eclosed female flies were put on 10 mL vials of standard cornmeal media for control experiments while 10-15 newly eclosed female flies were put on food containing all-trans-retinal (0.02% by weight retinal) for optogenetic experiments. All vials were wrapped with aluminum foil to prevent retinal degradation and to keep conditions similar in the control vials. After 3-5 days on the control food or 4-5 days on the retinal food, flies were starved by placing them in empty scintillation vials with half of a damp Kimwipe (20 ul of water/half wipe) for 15-21 hours prior to experiments. Experimental flies were anesthetized on ice prior to placing them into the behavioral arenas.

### Behavioral experiments

Behavioral experiments have been previously described in detail (Tao et al., 2020). In brief, experiments were conducted in a 4 cm radius circular arena with a 1.2 cm radius concentric light circle. Flies were given a 5-minute light acquisition period followed by a 10-minute dark acquisition period that reflected experimental conditions. The arena was lit with infrared light to enable tracking. The light circle was illuminated with red light (617 nm) for the last 3 minutes of each 6-minute experiment. The fly’s locomotion was recorded at 30 frames per second using an infrared video camera (Basler acA20400- 90umNIR). Recorded videos were compress to ufmf format before tracking (Branson et al., 2009). The tracking code models flies as an oval (using the MATLAB function regionprops) to extract the body orientation and centroid positions. Head position was tracked with the criterion that the current head position should be the endpoint along the major axis that makes the smaller turn from the previous head position.

### Kernel Density Estimates (KDE) of spatiotemporal distributions

For each genotype, each fly’s radial head position was aligned by first entry. To account for differences in first entry, all head positions after the 6-minute mark were set to nan and not considered. Spatiotemporal distributions of head position were then estimated using MATLAB’s ksdensity function with a gaussian kernel. First entry was defined as the first time the fly’s head enters the light zone (1.2 cm radius circle) after the light turns on.

### Electrophysiology

Single sensillum recording was performed using standard techniques (Schlief and Wilson, 2007). Orco- Gal4>UAS-Chrimson flies were held in a pipette tip using dental wax with the antenna accessible. The antenna was positioned using glass hooks and visualized using a microscope. A single sensillum was impaled with a glass pipette filled with saline. Responses were passed through a 100x amplifier and filtered with a 5 kHz low pass Bessel filter.

Flies were illuminated using a red (617 nm) light emitting diode (LED) (Thorlabs M617L3) connected with a LED driver (Thorlabs LEDD1B) with the intensity modulated by controlling the voltage using MATLAB. As with the behavior experiments, the LED light was collimated (Thorlabs ACL2520U) and focused using a plano-convex lens (Thorlabs LA1433). To deliver the same range of stimulus intensity in the electrophysiology experiments as the behavioral arena, we first measured the light intensity in the behavioral arena using a photometer (Thorlabs S121C) with a 1 mm diameter precision pinhole (Thorlabs P1000D). We then placed the LED at a distance from the fly such that the range of intensity values measured from the arena maps to voltage values between 0 and 5. To calibrate the light, we applied a series of voltage steps from 0 to 5 volts in intervals of 0.5 volts and measured the intensity using the precision pinhole. We then fit a shifted rectified linear function to map the voltage to intensity. Using these measured conversions, six 60-second behavioral positional trajectories were converted to from movement paths within the behavioral arena to light intensities that were exposed to flies during recording sessions (Figure 2-S1).

### Spike sorting and Spike Rate analysis

Data collection and spike sorting was conducted utilizing a custom MATLAB graphical user interface (GUI). The local field potential (LFP) was found by applying a 300-millisecond median filter to the signal. Then the raw voltage trace was baseline subtracted by subtracting out the LFP. Spikes were identified based on valleys in voltages below -0.8 mV with a minimum time lapse between spikes of 5 ms. Spikes were sorted based on shape and size by using principal component analysis followed by k-means clustering and then manual inspection. In this study, we recorded from ab1 and ab2 sensillum. Within the ab1 sensillum, there are 4 types of neurons: ab1A-D (de Bruyne et al., 2001; Larsson et al., 2004b). Meanwhile, within the ab2 sensillum, there are 2 types of neurons: ab2A-B. These neurons are differentiated by their waveform and spike amplitude. We only considered spikes from ab1A, ab1B, and ab2A neurons as these neurons showed a large change in firing rate during optogenetic stimulation. Spike rate was estimated using kernel smoothing with a 150 ms bandwidth (Shimazaki and Shinomoto, 2010).

### Filter analysis

To derive the linear filters, we delivered 6 sample patterns of 60-second trajectories of light intensity from our Orco dataset (Figure 2-S1). The patterns were up-sampled from 30 Hz to 10 kHz for single sensillum recordings. We recorded from 2 ab1 sensilla and 1 ab2 sensilla for each stimulus pattern. For each stimulus pattern, the trial with the least LFP baseline drift was baseline subtracted and set as a template. All other LFP for the stimulus pattern were linearly registered to the template LFP using a linear least squared rigid registration method (Umeyama, 1991). The average LFP was calculated from the registered LFP traces. The LFP, firing rate, and input stimulus were down-sampled to 100 Hz for filter analysis.

The firing rate was modeled as a two-stage linear-linear filter cascade. The linear filters were fit based on previous methods (Pillow et al., 2008). Taking the first transformation from input stimulus into local field potential as an example, we calculated each linear filter as follows. First, given a *T* time point stimulus train with (*d* − 1) time point zero padding at the beginning, we can generate a stimulus matrix *S* ∈ ℝ^*Txd*^ with feature length *d* Each row that corresponds to a time point *T* contains the stimulus train from time *t* − *d* to *t*. This is equivalent to a hankel matrix of the zero padded stimulus train. Next, we can define the response vector *r* ∈ ℝ^*Tx*1^ from the average local field potential from single sensillum recordings using the stimulus train. From the stimulus matrix and the response vector, we can pose the stimulus to response as a simple linear regression with a linear filter k:

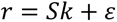

We solved to *k* using Tikhonov regularization, minimizing

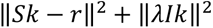

The regularization parameter *λ*, which helps to prevent overfitting, was chosen based on the elbow point in the log-log plot of the regularized solution norm and the residual norm (Figure 2-S1B) (Lawson and Hanson, 1995). The linear filter from LFP to firing rate was calculated using the same method.

### Quantification of kinematics

Speed and curvature were calculated as described previously. Briefly, given the two-consecutive center of mass positions (*p*_1_, *p*_2_), we can calculate speed as

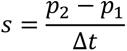

Then defining curvature (*k*) as the change in the vector (*N*) normal to the movement path, we get:

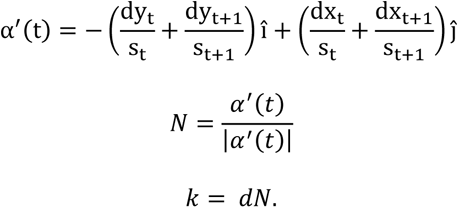

### Behavior quantifications

Behaviors were characterized using three different methods:

1. KDE of spatialtemporal distributions as described above (Figure 1)
2. Radial occupancy: The overall probability mass distribution of the average fly being a radial distance away. A bin size of 2 mm (0.05 radial units) was utilized in generating the distribution. (Figure 1-S1 and Figure 7)
3. Radial density of turns: This is the the probability mass function of the density of head positions during the middle of sharp turns (see below). A bin size of 2 mm (0.05 radial units) was utilized in generating the distribution. (Figure 7)
4. Probability of being inside: This is the proportion of flies inside the central 1.2 cm radial location as a function of time. A 200 ms mean filter was used to smooth the trace. (Figure 1-S2)

### Definition of movement states

Each fly’s movement path was classified into one of four locomotor states (stop, boundary, sharp turn, curved walk) based on a previously described method (Tao et al., 2020). Briefly, stops occurred when the speed was less than 0.5 mm/s. Flies were in a boundary state when the center of mass was within 1.5 mm (half a fly length) of the arena boundary. Sharp turns occurred around large peaks in curvature while curved walks did not contain large peaks in curvature. A set of 1-2 kinematic features and duration defined each locomotor state. Moving forward, we will group duration into kinematics in the terminology for brevity. Stops were characterized by the total curvature (reorientation) and duration. Boundary states were characterized by the total angle of the arc of movement around the boundary and duration. Sharp turns were characterized by the total curvature, average speed, and duration. Finally, curved walks were characterized by the average curvature, average speed, and duration.

### Parameterization of movement states based on kinematics

We separated the boundary state trajectories into before and after first entry. First entry was defined as the first time the ORN firing rate exceeded 10 Hz. The total angle of the arc of movement around the boundary and duration was fit to a bivariate lognormal distribution. All other state kinematic distributions were generated based on ORN activity. Taking sharp turns as an example, we calculated the mean firing and mean change in firing for the 200 ms interval leading up to the initiation of each sharp turn instance.

Sharp turns were then separated into three categories based on the calculated ORN activity history: These were before first entry, after first entry with baseline firing rate, and after first entry with a non-baseline firing rate. For sharp turn trajectories before first entry, we independently fit each kinematic feature to a lognormal distribution. For sharp turn trajectories with baseline firing rate after first entry, we fit the kinematic features of the first two trajectories after reaching baseline and trajectories 3 and later with two separate independent lognormal distributions. These trajectories were separated since the kinematics of the first two trajectories after reaching baseline tend to exhibit large differences from trajectories before first entry (Figure 4-S4). However, later trajectories tend to display similar kinematics as trajectories prior to first entry. We used the Wilcoxon rank sum and estimation methods to test for significant changes in baseline firing rate kinematics from before first entry (Ho et al., 2019). In Figure 4-S4B, scatter plots show individual data points and corresponding error bars show mean and bootstrapped 95% confidence interval (resampled 10000 times, bias-corrected, and accelerated). 95% confidence interval for differences between means were calculated using the same boostrapping methods.

For sharp turn trajectories with non-baseline firing rates after first entry, we generated a kinematic mapping based on neural response that is described in the next section below. Since this mapping does not take into account kinematic changes over prolonged periods of inhibition in ORN activity after the fly leaves the light ring, we fit time dependent lognormal, beta, and exponential distributions to speed, curvature, and duration respectively. To do this, we first implemented a sliding window of 0.5 seconds with a 0.3 second overlap over the time since the start of each inhibition period. We then fit the appropriate distribution (i.e. lognormal for speed) over the kinematics of the sharp turn trajectories that started within the time window. We then interpolated the distribution parameters over time using a spline function. We repeated this process for curved walks and stops. The distribution fits for after first entry with baseline firing rate and after first entry with inhibition firing rate were used in the agent-based model described in a later section

### Neural Response to Kinematic mappings

We used a K-nearest neighbors (KNN) approach to generate kinematic mappings for each state based on neural responses. The goal is to estimate the distribution of average kinematics given recent ORN activity. Using curved walk speed as an example, we first calculated the average firing rate (*f*) and change in firing rate (Δ*f*) for the 200 ms window prior to the start each curved walk trajectory. This allowed us to embed each curved walk as a point in the (*f*, Δ*f*) space (Figure 4-S1A).

To obtain the distribution of potential future curved walk speeds for a given (*f*, Δ*f*) coordinate, we first divided the (*f*, Δ*f*) space into grids defined by the intersection of Δ*f* spanning from -150 spikes/s^2^ to 150 spikes/s^2^ in 15 spikes/s^2^ increments and *f* spanning from 0 spikes/s to 55 spikes/s in 1 spikes/s increments. The range of coordinates in the (*f*, Δ*f*) space was chosen to span over 99% of the possible state points within the dataset. At each coordinate in this grid, we want to use the K closest points – in terms of Euclidean distance – to compute a distribution of curved walk speeds.

Prior to calculating the Euclidean distance between the coordinate and all curved walk points in the (*f*, Δ*f*) space, we first applied a weight to each point. In this study, we weighted the *f* and Δ*f* of each point by dividing by 10 and 30 respectively. These weights – which are selected heuristically – were selected since the maximum of the absolute value of the Δ*f* is ∼3 times more than the maximum of the *f* of all of the state points. It should be noted that in retrospect, only the relative weights matter since a factor of 1 and 3 for *f* and Δ*f* will be equivalent as the weights of 10 and 30. Another reason we chose to have a higher weight along the Δ*f* axis is because we expect that two trajectories that differ by 1 spikes/s in firing rate will have more similar locomotor kinematics than two trajectories that differ by 1 spikes/s^2^ ORN firing rate. Furthermore, the change in firing rate is noisier when compared to the firing rate since Δ*f* is calculated by multiplying the difference in firing rate by the frame rate (30 Hz).

Since there are locations in the (*f*, Δ*f*) space where there are little to no data points, we defined a maximum Euclidean distance bound (T) that points have to fall within to be considered as part of the distribution. Using this bound prevents curved walks trajectories where flies are experiencing extremely different ORN activity from influencing the distribution for a grid coordinate. This means that we can represent the Euclidean distance bound (T) as an ellipse:

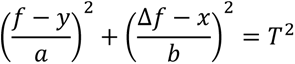

Where *r*, *S* are the weights for *f* and Δ*f* respectively and *y*, *x* are the coordinate locations within the (*f*, Δ*f*) space. To summarize, for each *y* and *x* coordinate location in the (*f*, Δ*f*) space, we are fitting the curved walk speed values of the K closest points – that fall within an ellipse centered at the coordinate location – to a lognormal distribution (Figure 4-S1A). Lognormal distributions were only fit for coordinates with more than 15 trajectories (points) within bounds as a low sample size will lead to inaccurate estimates of the underlying distribution. After fitting lognormal distributions to each coordinate location within the (*f*, Δ*f*) space, we performed linear interpolation of the lognormal parameters to get the distributions of curved walk speed any arbitrary *f* and Δ*f* location in the space.

The values of K and T were selected by first calculating the standard error of the mean (SEM) over a grid search of K and T and then choosing a value near the elbow point (Figure 4-S1C). Based on this criterion, sharp turn and curved walk kinematics were mapped to the neural response space using a K of 64 trajectories and a T of 1.5. Stop kinematics were mapped to the neural response space using a K of 64 trajectories and a T of 1.

### Adaptation in Neural Response to Kinematic mappings

To calculate the kinematic mappings in the (*f*, Δ*f*) space as a function of time since first stimulus experience, we extended the KNN method by introducing time since first entry (*t*) as a third dimension. Here, we divided the (*f*, Δ*f*, *t*) space into grids defined by using the same grid interval in the *f* and Δ*f* directions. In the time dimension, we used grids spanning from 0 to 180 seconds (duration of the light on period) in 5 second increments. In this 3-dimensional space, the Euclidean bound (T) becomes:

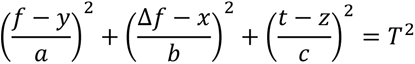

Where *r*, *S*, *T* are the weights for *f*, Δ*f*, and *t* respectively and *y*, *x*, *z* are the coordinate locations within the (*f*, Δ*f*, *t*) space. To summarize, for each *y*, *x*, and *z* coordinate location in the (*f*, Δ*f*, *t*) space, we are fitting the curved kinematic values of the K closest points – that fall within an ellipse centered at the coordinate location – to a lognormal distribution (Figure 4-S1B). In this study, we used weights of 10, 30, and 20 respectively based on the same reasoning described in the previous section. We used the same K and T as the (*f*, Δ*f*) space kinematic mapping (see previous section).

### Neural Response to turn optimality

Turn optimality ratio was defined as the probability that a fly will turn in the direction that will take the least amount of turning to re-orient itself towards the center of the light ring. To calculate the turn optimality, we define the current movement direction as a vector (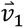) starting at the center of mass position 200 ms prior to the state transition (p_1_) and ending in the center of mass position at the state transition (p_2_).

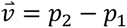

Next, we defined a vector that points radially inwards towards the center of the arena from the sharp turn index (*p*_2_) as:

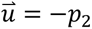

Then we defined a vector normal to the xy plane (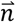). From this, we calculated the directed angle of the current direction relative to the inward vector.

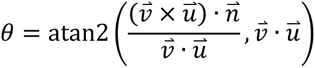

When the directed angle is positive, then turning left is optimal. Meanwhile, a negative directed angle indicates that rightward turning is optimal. Since a positive curvature represents a leftward turn and a negative curvature represents a rightward turn, a fly makes an optimal turn if the sign of the total sharp turn (or stops) curvature or average curved walk curvature during the next state instance is the same as the sign of the directed angle. The turn optimality ratio for a given state was defined as the total number of optimal state trajectories over the total number of state trajectories. Turn optimality was mapped to the neural response space in the method described above using a K of 64 points and a T of 1.5.

### Neural response to transition probability

Transition probability was defined as the number of non-self-transitions from one state to another state. This was mapped to the neural response space in the method described above using a K of 128 points and a T of 1.5. After mapping to the set of locations *Y*_*i*_ in the ORN activity space, we implemented a 5x5 (75 Hz/s^2^ x 5 Hz) convolutional filter in order to smooth out noise due to low sample sizes especially in regions with high *f* and Δ*f*. Finally, all other locations in the ORN activity space were found using linear interpolation. The transition probabilities are described below and can be generated using the accompanying code. Sharp turns always transitioned to curved walks. Curved walks largely transitioned to sharp turns except when the firing rate is low, where there is an increase in transition to stops. ∼25 percent of stops transitioned to curved sharp turns and the remainder of stops transitioned to curved walks. This transition probability did not show any noticeable trends in the neural response space and is largely similar across genotypes.

### Modeling ORN rules of summation

The goal is to understand the combinatorial effect of activating different ORN classes on changes in sharp turn and curved walk kinematics during odor-guided locomotion. To do this, we first divided our ORN activity space into five distinct regions (Figure 2C and Figure 6-S1A). Region I consist of large positive increase in ORN firing rate defined by a threshold of 20 Hz/s^2^. Region II consists of high firing rate defined by a threshold of 15 Hz. Region III consists of large negative decrease in ORN firing rate as defined by a threshold of -20 Hz/s^2^. Region IV consists of an inhibition of firing rate as defined by a threshold of the baseline 4.7 Hz firing rate. Finally, region V consists of low ORN firing rate between baseline and 15 Hz. Within each region, the state kinematics followed an approximately lognormal distribution (Figure 6-S1A). We chose to use these large regions relative to the previously state method of kinematic mapping to neural response because:

1.) The larger number of samples in these broader regions reduced sampling error

2.) Kinematic changes were generally consistent within each of these regions for the same genotype (Figure 4 and Figure 4-S2).

Lognormal distributions are parameterized by the mean and variances of the log of the state kinematics (Figure 6-S1A). We next found that activation of combinations of ORN classes did not result in a direct sum of means when activating individual ORN classes within the combination and that increasing the number of ORN classes activated did not lead to a monotonically change in the effect on the mean (data not shown). This suggested that two different ORNs have an overlapping influence on kinematics and this overlap is distinct based on ORN identity. These observations were even more pronounced when looking at the variances, which suggests that two different ORNs have a correlated effect on locomotor parameters.

To model these effects, we considered a simple linear regression model for the means and variances of the lognormal distributions that describe the effect of ORN activity in each of the 5 regions on state kinematics (Figure 6-S1B and 2). Using sharp turn curvature in region 1 as an example, in this formulation, the mean and standard deviation of the log of the sharp turn curvature in region 1 when activating a single class or group (i.e. Orco) of ORN is:

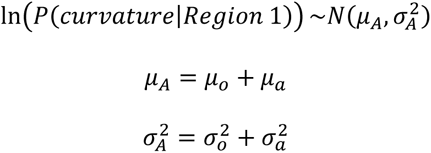

Extending this to the simultaneous activation of a set of two classes of ORNs (A and B) is formulated as:

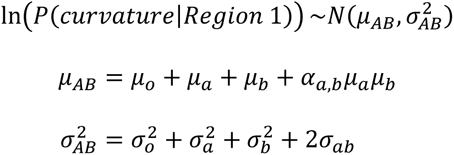

Where *μ*_o_ and 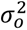 are the genotype specific baseline mean and standard deviation calculated from before first entry. *μ*_a_, *μ*_b_ and 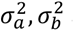 are the influence of the ORN class A and B on the mean and standard deviation of the state kinematics. Overlapping influences of ORN class *a* and *b* is modeled using the coefficient (α_*ab*_). Finally, *σ*_*ab*_ is the covariance between classes a and b. The model was fit using MATLAB’s maximum likelihood estimation function. We fit a total of 7 ORN combinations: Orco + Ir8a, Ir64a + Ir75a, Or42b + Or92a, Or42b + Ir64a, Or42a + Or42b;Or92a, Or42b + Ir64a;Ir75a, and Ir75a + Ir64a;Or42b for sharp turn average speed, sharp turn total curvature, curved walk average speed, and curved walk average curvature. In this paper, we show results for the four combinations involving Or42b (Figure 6).

### Interpretation of the model in the kinematics space

We can transform the lognormal means into the real-world state kinematics space by taking the exponent:

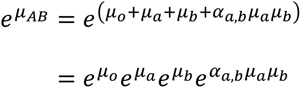

We note that the influence of each genotype (*e*^*μ_a_*^, *e*^*μ_b_*^) and the full interaction term (*e*^*α_a,b_μ_a_μ_b_*^) each act as a multiplier to influence the real-world state kinematics. We can define these terms as the gain in kinematics due to each genotype and their interactions respectively.

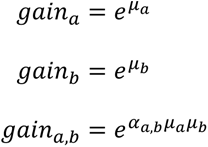

A gain of less than 1 indicates a reduction in kinematics (i.e. decrease in speed) while a gain of greater than 1 indicates an increase in kinematics. Here, we set a threshold of 0.1 to indicate whether the gain caused by a genotype or interactions between genotypes leads to a notable change in kinematics. This means that:

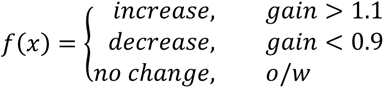

### Definition of synergy and antagonism, dominance, and other interactions

Independent activity of single classes or groups of ORNs can cause increases (*S*_*r*_ > 0) or decreases (*S*_*r*_ < 0) in kinematics. When two separate classes or groups of ORNs are co-activated using Chrimson, *α_a,b_μ_a_μ_b_* captures the potential effect of convergent downstream interactions that influence locomotor kinematics (Figure 6-S2A). These interactions are defined to be synergistic if the interaction acts to enhance the individual effects of single ORN class effects and the interaction results in a notable change in kinematics (see above section). The interactions are defined to be antagonistic if the interactions act to cut back on the individual effects of single ORN class effects and the interaction results in a notable change in kinematics (see above section). For instance, if independent activity of single classes of ORNs both cause a decrease in kinematics, then the interaction will be synergistic or antagonistic if the interactions cause a decrease in kinematics or increase in kinematics respectively. Cases where the individual ORN classes cause opposing effects will result in other effects that cannot be directly classified as synergistic or antagonistic (Figure 6-S2B). In these cases, one possibility is that the resultant change in kinematics may be dominated by the activation of a single class of ORN. For instance, if activating ORN class a cause an increase in kinematics, ORN class a cause a decrease in kinematics, and activating both ORN a and b causes an increase in kinematics, then the change in kinematics will be dominated by ORN class a. Alternatively, there are cases where both ORNs do not cause a change in locomotor kinematics on their own but activating both causes a notable increase or decrease in in kinematics. These cases are labeled as other interactions. Finally, when the interaction term does not cause a notable change to kinematics, then the two ORN groups likely do not interact and will sum linearly through parallel pathways.

### Connectomics analysis

To determine whether there are connections between ORNs and LHONs we first queried the Hemibrain connectomics database to extract all input and output connections for the ORNs of interest, and all identified classes of uniglomerular PNs (uPNs) and multiglomerular PNs (mPNs) (Scheffer et al., 2020). We then identified which of these PN classes made strong (>9) connections to any of the LHONs characterized in a recent study (Dolan et al., 2019). Because only the right hemisphere in Hemibrain is complete, any left hemisphere connections were excluded due to the possibility that connection values would be unreliable. Based on the connections between ORNs and uPNs/ mPNs and between uPNs/ mPNs and the LHONs we characterized which LHONs receive inputs from ORNs either directly via their cognate uPNs or indirectly via mPNs that receive input directly from the ORNs or from their cognate uPNs (Figure 6-S3).

### Agent-based model

Virtual flies were initialized as described previously (Tao et al., 2020). Each simulation was run using 250 flies modeled as point objects for 6 minutes with the center light ring turning on at the 3-minute mark to match experimental protocol. The simulations were run at 100 Hz since the stimulus to ORN firing rate filters were computed at 100 Hz. After the 3-minute light off period, the firing rate of flies were calculated using the previously derived linear filters and based on the light stimulus experience as a function of radial distance from the center. First entry was defined as the first time the calculated ORN firing rate is above 10 Hz. Before first entry, locomotor kinematics, turn optimality, and state transitions was calculated based on previously described distributions calculated from empirical flies before first entry. After first entry, state transitions, locomotor kinematics, and turn optimality were sampled based on kinematic mappings in the neural response space. When ORN activity is at zero (inhibition), the locomotor kinematics were sampled from inhibition distributions based on the time since the start of the inhibition period. Virtual flies performed sharp turns by moving straight for half of the duration of the sharp turn at the sampled speed, then turning based on the sampled curvature over the course of one-time step, and finally, moving straight for the remainder of the time step. Stops were implemented in the same manner, except that the speed is set to zero. Virtual flies performed curved walks by moving at the average speed and curvature for the duration of the curved walk. When the fly reaches within 1.5 mm (0.0375 normalized distance) of the arena boundary, the virtual fly enters the boundary state. Here, the fly moves around the arena boundary at a constant angular speed and duration sampled from the empirical before and after first entry distributions. Flies exit out of the boundary state by reorientating towards the center of the arena and selecting a curved walk or sharp turn state. From the set of 250 virtual flies, only the flies with first entry times within the 85^th^ percentile of empirical first entry times were kept. For Figure 7C, border choice was implemented for curved walks by imposing an exponentially time decaying probability of state transition after reaching a threshold of +/-15 Hz/s^2^.

### Correlation analysis for turn density and radial occupancy

Synthetic fly radial occupancy was subtracted from empirical fly radial occupancy to obtain the difference in radial occupancy. Since the sum over radial distance for the difference in the probability mass function equals zero, the total positive difference measures the level of discrepancy between the empirical and synthetic flies (Figure 7-S4A/C). The following is repeated for radial density of turns. Genotypes, where there is a larger positive position or turn difference, has a larger correlation between the difference in radial occupancy and turn density (Figure 7-S4C).

## Acknowledgment

We would like to acknowledge the members of Bhandawat lab for discussions and Barani Raman for carefully reading the manuscript. This research was supported by RO1DC015827 (VB), RO1NS097881 (VB) and an NSF CAREER award (IOS-1652647 to VB), and an NIH F31NS120835-02 (LT).

## Author contributions

VB: Conceptualization; Supervision; Funding acquisition; Writing;analysis. LT: Conceptualization, Experimentation, analysis, Writing. SPW: Experimentation, analysis, Writing;

---

**Figure 1S1.**
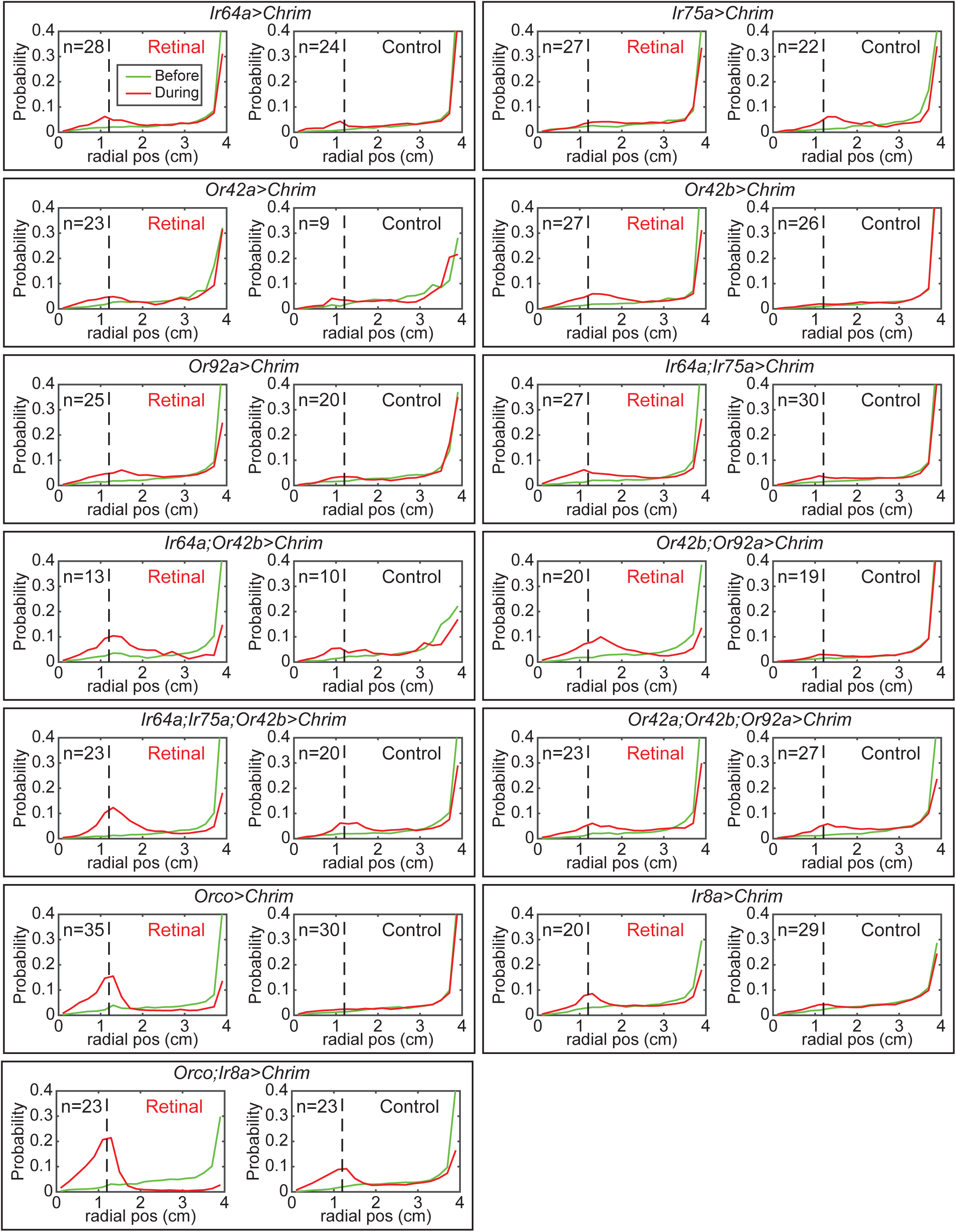
Radial occupancy of flies under optogenetic stimulation of different subsets of ORN classes. Radial occupancy for before first entry is shown in red and after first entry is shown in green. Dotted lines show nominal light border (1.2 cm). Control flies were not fed retinal, which is necessary for optogenetic activation of neurons.

**Figure 1S2.**
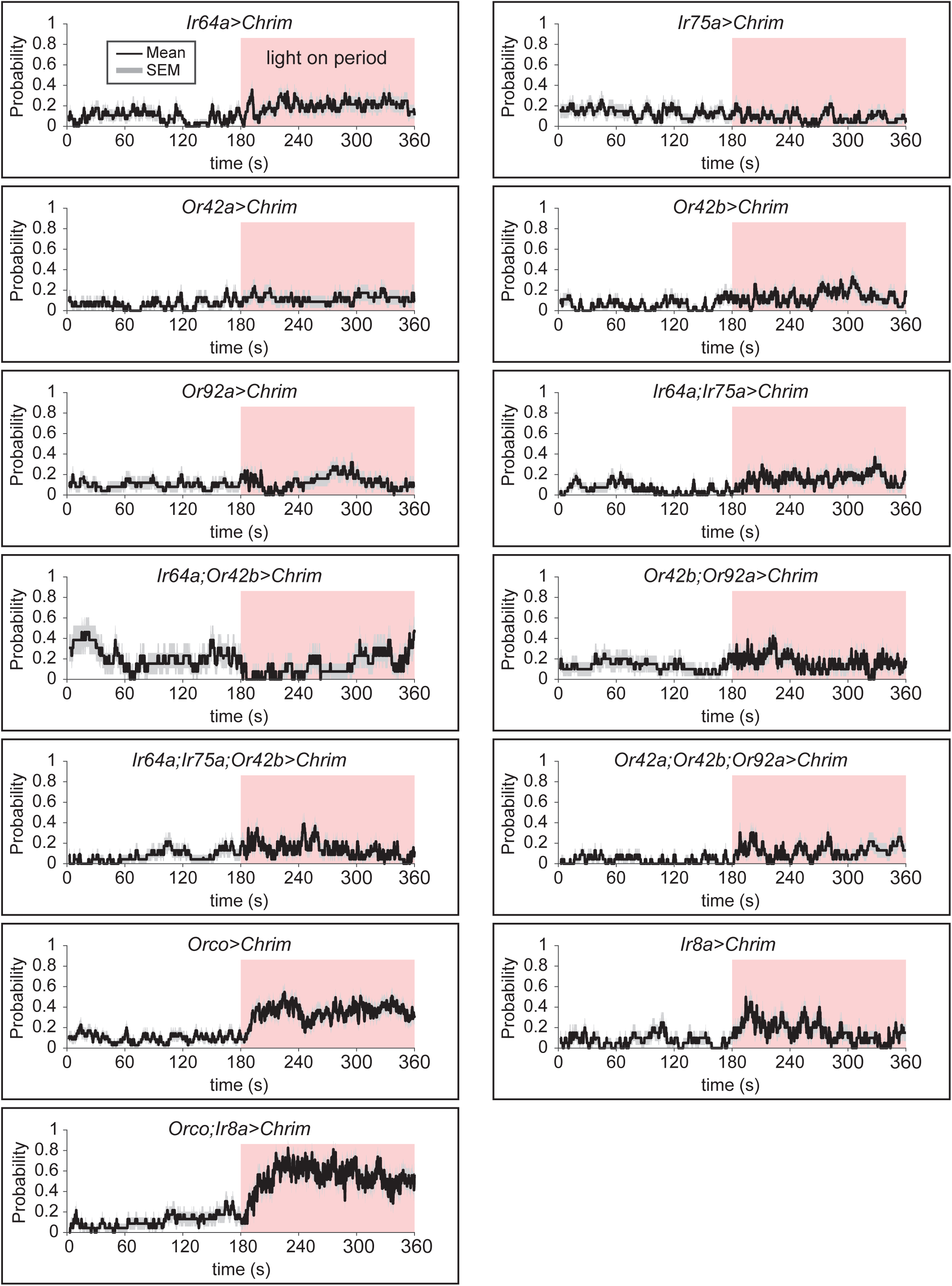
Proportion (probability) of flies located in the light zone under optogenetic stimulation of different subsets of ORN classes. The nominal light border is 1.2 cm. Light turns on at the 3 minute mark (180 seconds).

**Figure 2S1.**
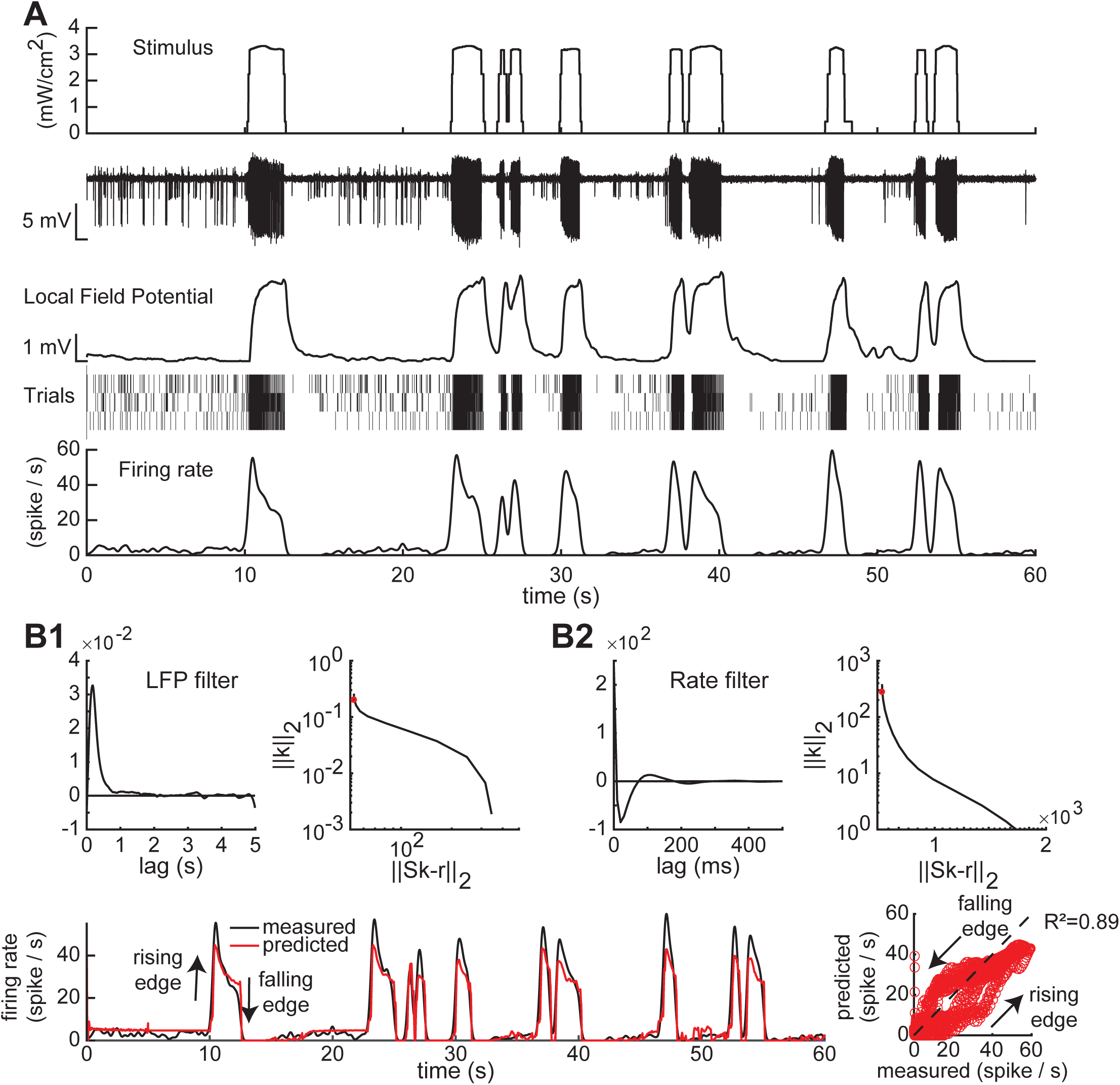
ORN responses, construction and validation of encoder. **A.** From the top to the bottom: A sample 60 second light stimulus; local field potential subtracted trace to show spikes; median filtered trace showing local field potential; Raster plot of 3 sensilla recordings for the stimulus pattern; the average firing rate profile over the 3 trials was estimated using a 150 ms gaussian kernel. **B1.** A linear filter was calculated using Tikhonov regularization (ǁSk-rǁ²+ǁλIkǁ²) to transform from the stimulus to the local field potential. **B1 Right.** The H-curve with the regularization parameter (red dot) corresponding to the LFP filter. **B2.** Same as **B1**, but for the linear filter transformation from the local field potential to the firing rate. **C. Left.** Measured (black) and filter predicted firing rates (red dot) for this stimulus pattern. The filter predictions underestimate the peak responses. **Right.** Plot of the measured and predicted against each other. The dotted line is the identity line. In general, the filter predictions is highly correlated to the measured, but underestimate the rising edge and overestimate the falling edge.

**Figure 2S2.**
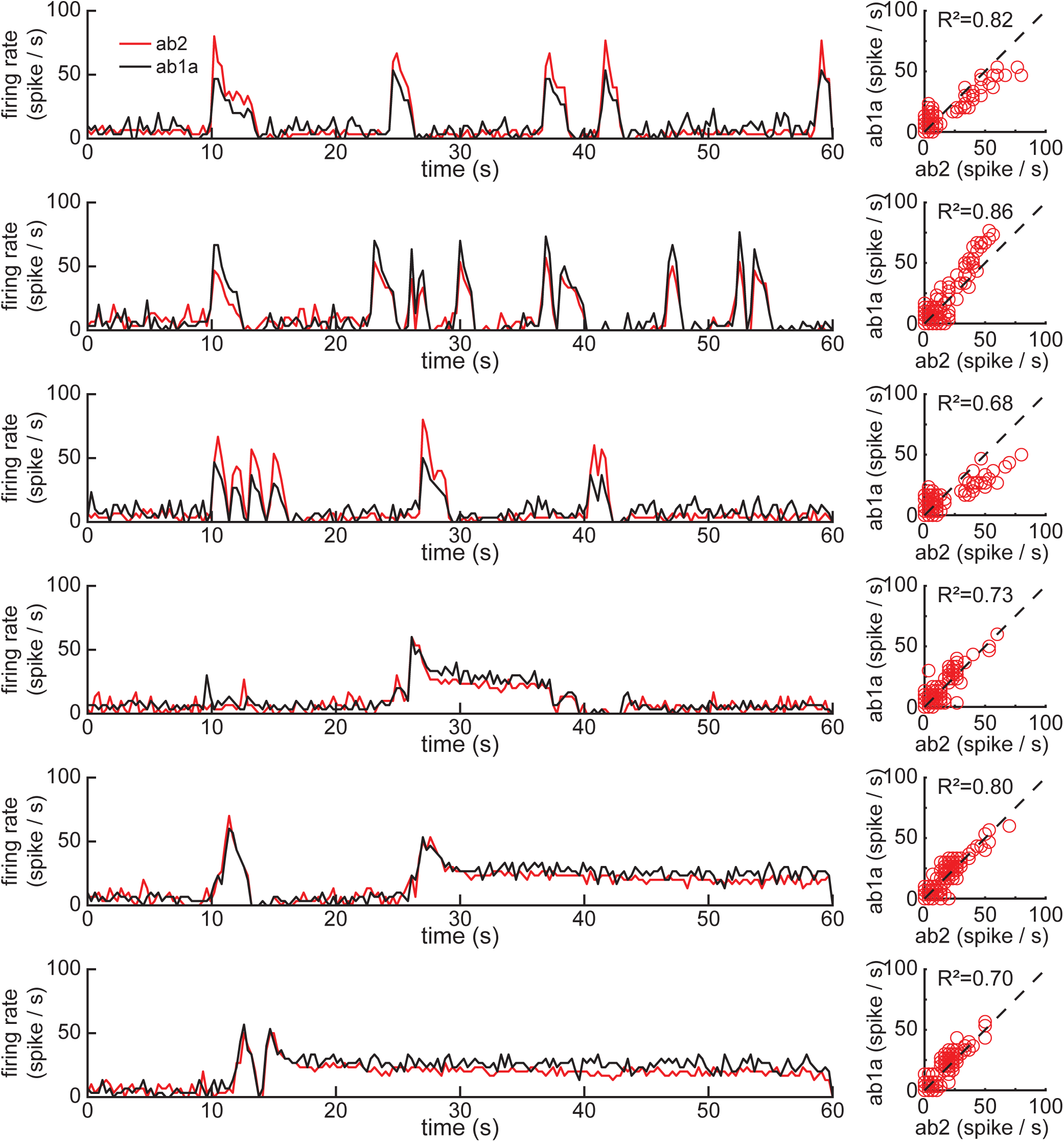
Optogenetic activation of ORNs trigger similar firing rate responses across ab1a and ab2 sensilla. Left. Each row shows firing histogram for a single trial of recordings from ab2 and ab1a sensilla in response to six types of stimulus pattern. **Right.** Scatter plot of the ab2 and ab1a histogram traces against each other. The dotted line is the identity line.

**Figure 4S1.**
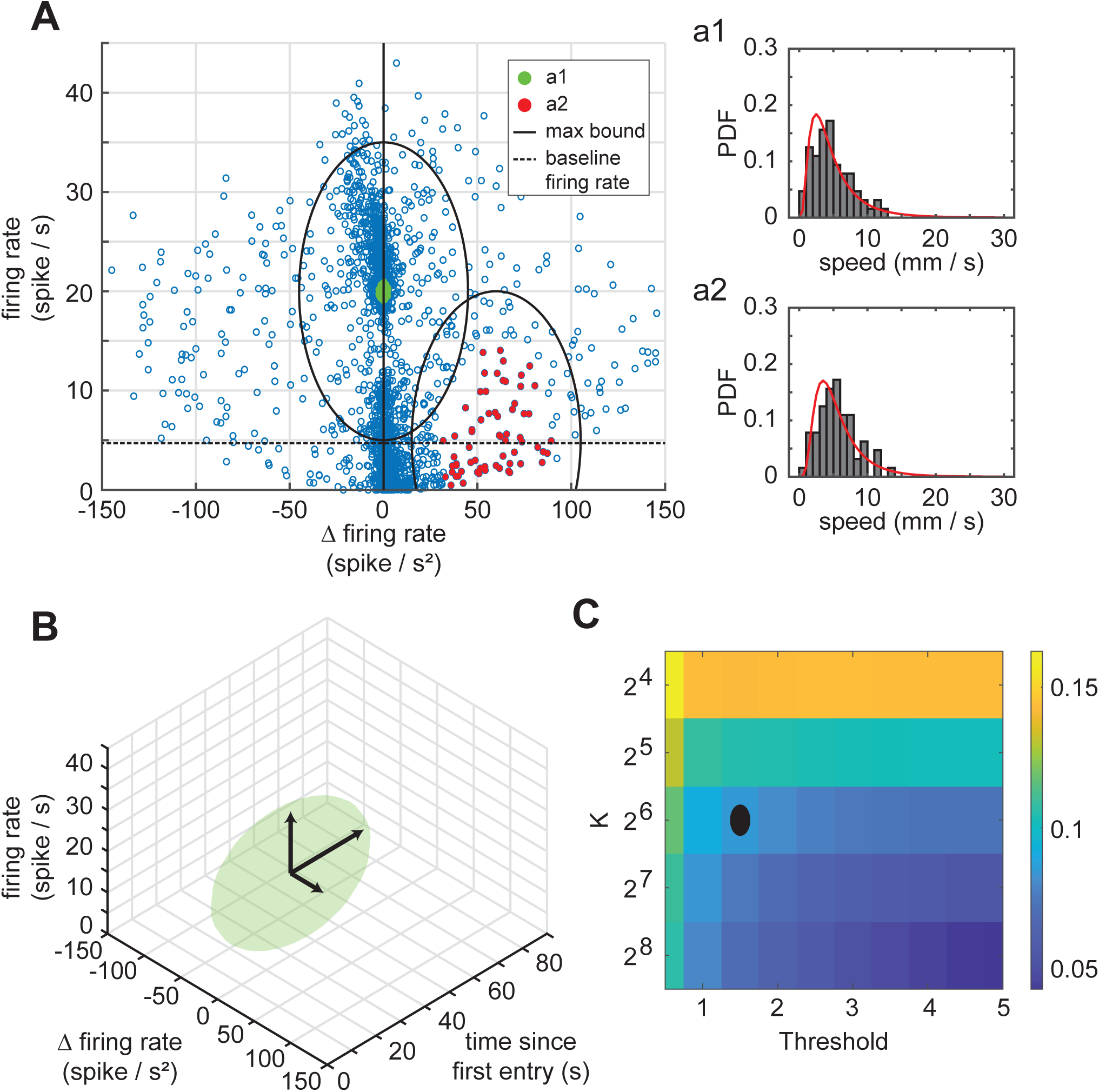
K-nearest neighbor approach to estimating how ORN activity drives locomotor kinematics. KNN approach for the estimation of speed of the curved walk. **A.** Each blue point is the average firing rate and change in firing rate of ORNs 200 ms prior to the start of a curved walk. Therefore, each blue point is associated with a curved walk that is just about to start and therefore a single speed value reflecting the averge speed of that curved walk. At each location in this grid, the K nearest trajectories are used to estimate a log-normal probability density function (PDF). **a1** and **a2** shows example log-normal PDF fits of curved walk speed corresponding to the green dots around 20 spikes/s firing rate, 0 spikes/s² change in firing rate and red dots around 5 spikes/s firing rate, 60 spikes/s² change in firing rate respectively. Dotted black line shows the baseline firing rate. Black solid line indicates the set maximum bounding distance. **B.** The maximum bound of **a1** extended to account for changes in ORN effect on locomotion due to adaptation. **C.** K and the maximum bounding oval threshold is determined by finding the inflection point (black dot) in minimizing the standard error of the mean across all grid points in the 3D space.

**Figure 4-S2.**
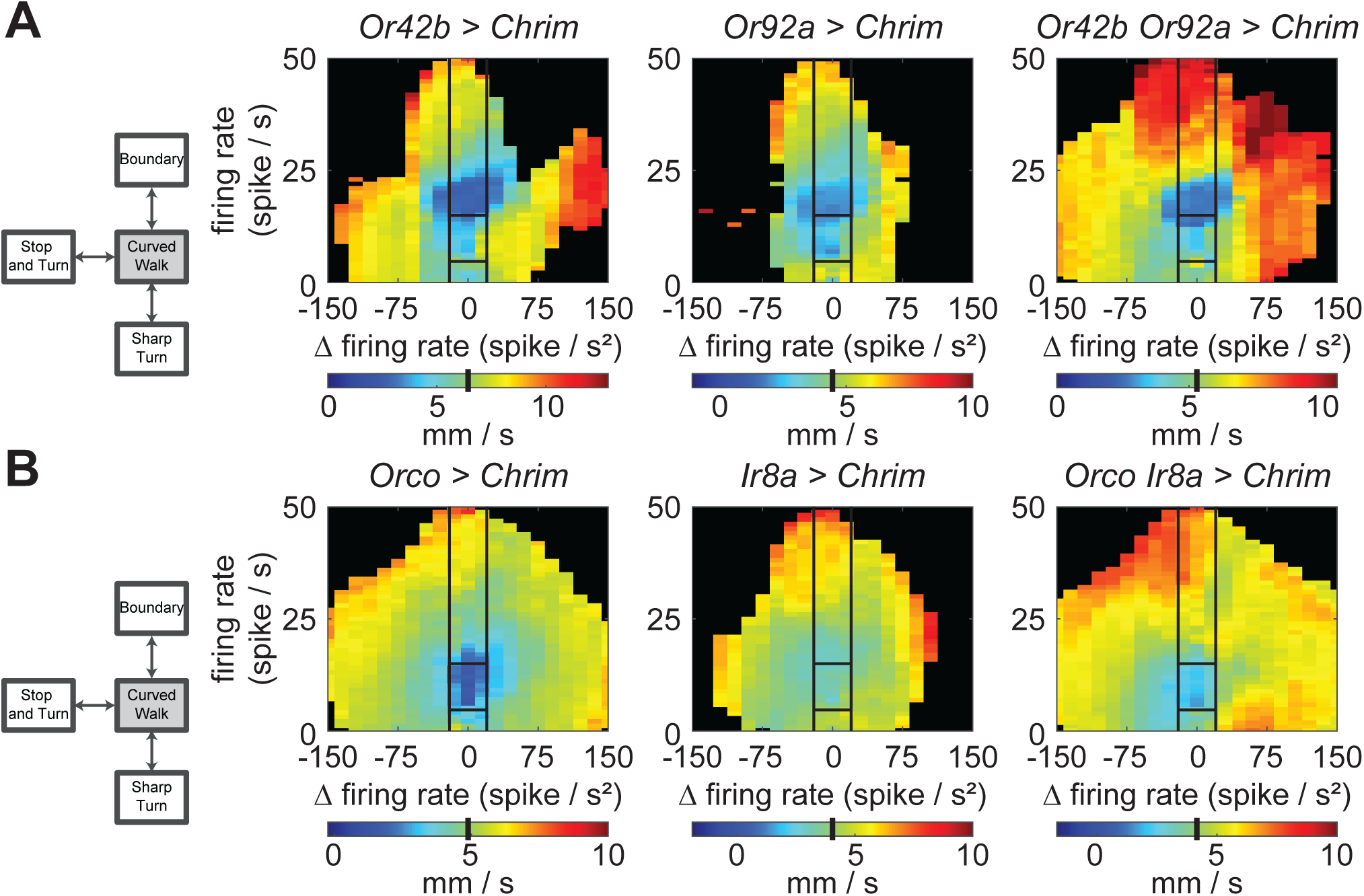
Some locomotor parameters are more affected when a small number of ORN classes are active. **A.** Curved walk speed is strongly affected by activation of a single ORN class (*Or42b* and *Or92a*). The effect is even larger when the two are activated together. Black lines separate out the sensorimotor mapping into 5 broad regions based on firing rate and change in firing rate. Baseline speed is shown as a black bar in the colormap. **B.** The effect of activated large populations of neurons that labels the co-receptor *Orco*, *Ir8a*, or either is smaller than that of individual ORNs.

**Figure 4S3.**
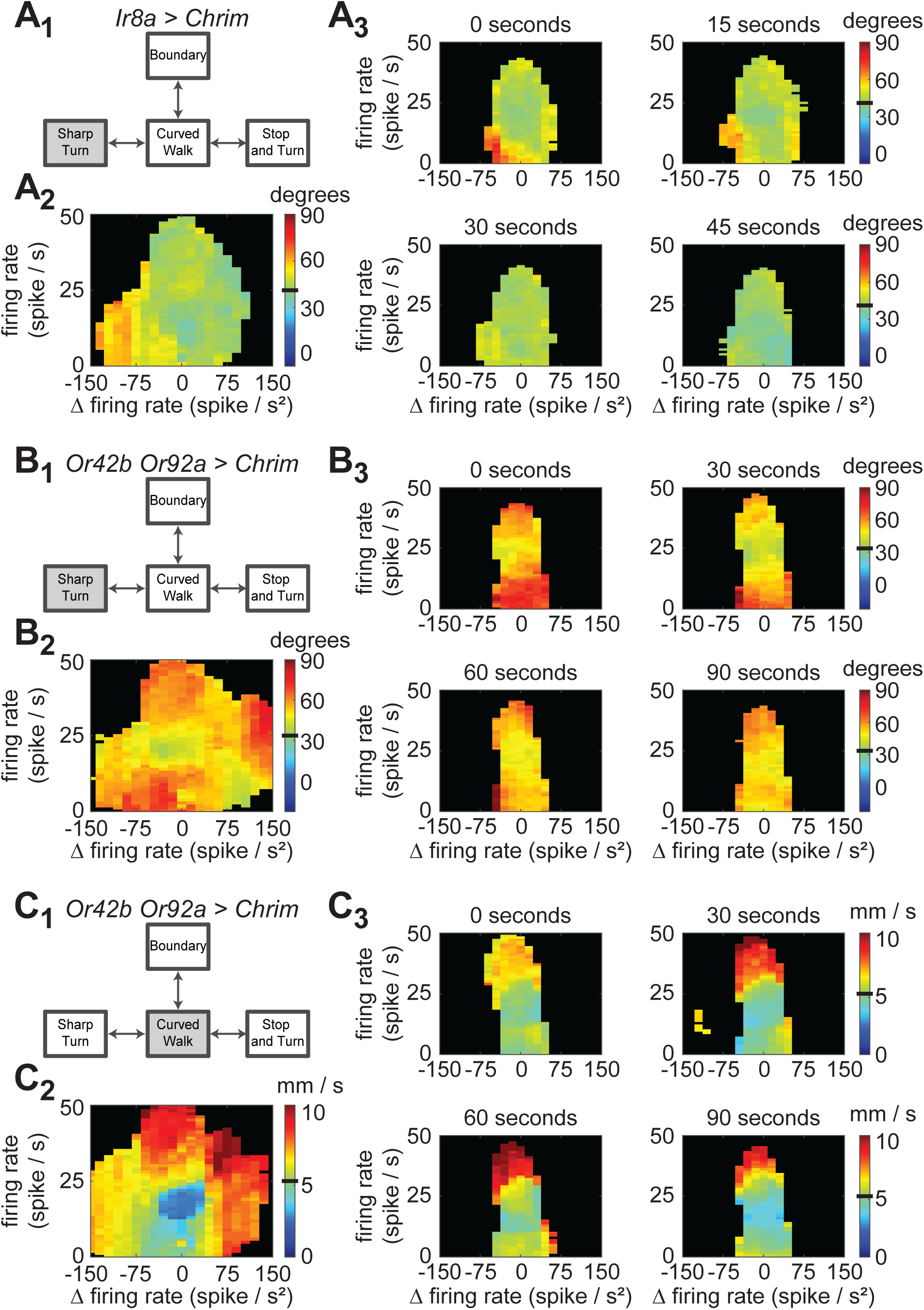
The effect of ORN activity on locomotor kinematics changes over time after first entry. **A.** KNN estimate of sharp turn curvature for *Ir8a* > Chrim (**A1**) across all time (**A2**). **A3.** KNN mapping of ORN activity to sharp turn total curvature showing the higher curvature due to a sharp drop in ORN activity only occurs within the first 30 seconds after first entry. **B.** KNN estimate of sharp turn curvature for *Or42b* and *Or92a* > Chrim (**B1**) across all time (**B2**). **B3.** KNN mapping of ORN activity to sharp turn curvature showing the higher curvature at high firing rates and during inhibition adapts, but is still present after 90 seconds since first entry. **C.** KNN estimate of curved walk average speed for *Or42b* and *Or92a* > Chrim (**C1**) across all time (**C2**). **C3.** KNN mapping of ORN activity to curved walk speed shows a large increase in speed within the first 30 seconds since first entry that does not continue to adapt.

**Figure 4S4.**
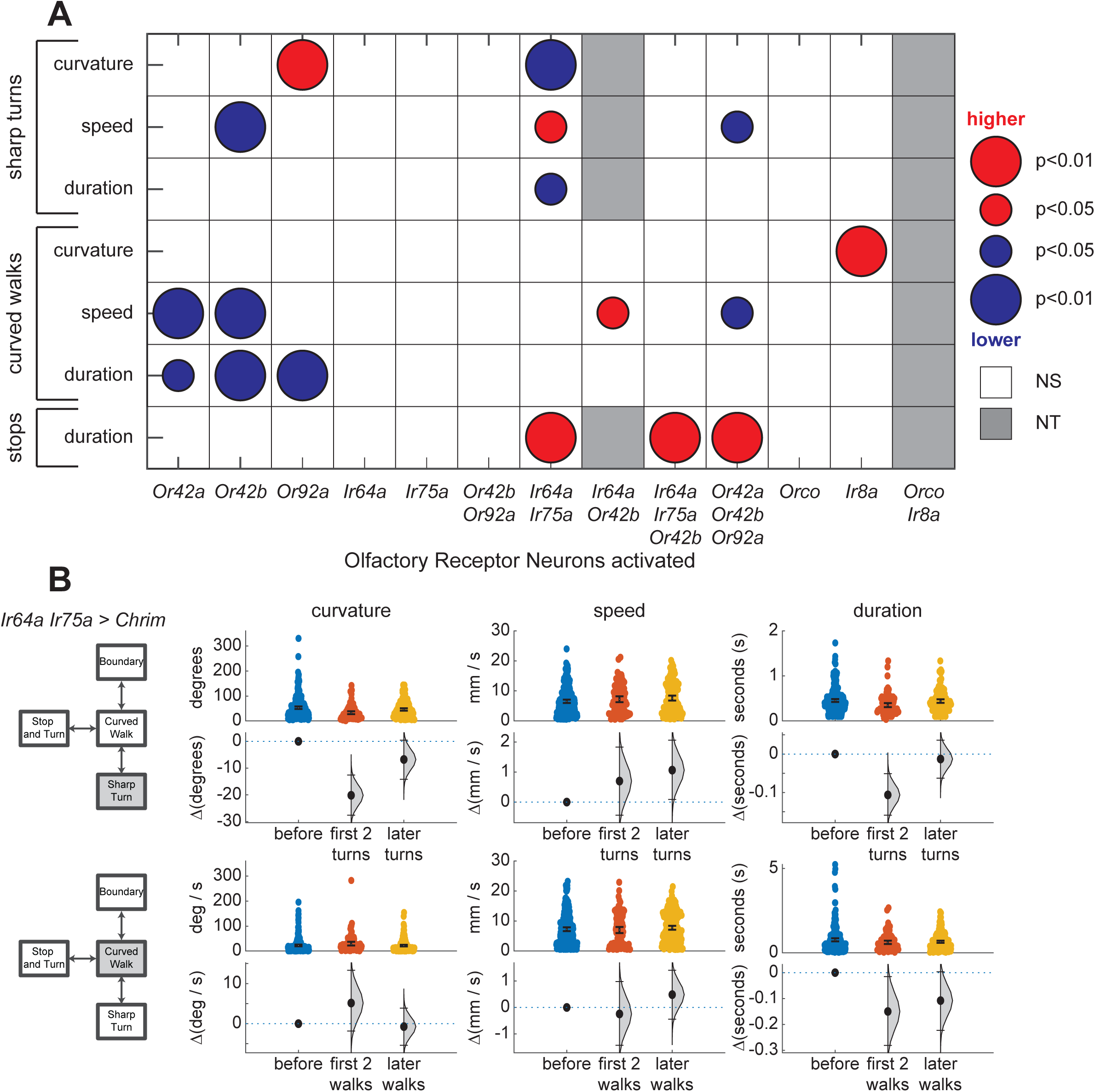
Previous ORN experience has limited effect on locomotion after ORN activity reaches baseline. **A.** One sided Wilcoxon rank sum tests comparing the sharp turn, curved walk kinematics and stop duration when the fly loses the odor source (ORN activity reaches baseline firing rate) is higher than (red) or lower than (blue) baseline values before first entering the light ring. Circle size shows significance at 0.05 and 0.01. Parameters that were not significant (NS) are in white. Parameters that had less than 30 samples were not tested (NT). **B.** For flies where ORN experience does have an effect on baseline kinematics, this kinematics adapts over time. *Ir64a* and *Ir75a* > Chrim retinal flies perform less curved sharp turns for lower time durations after recently reaching baseline firing. After spending more time at baseline, the sharp turn curvature and duration adapts to baseline levels before first entry.

**Figure 6S1.**
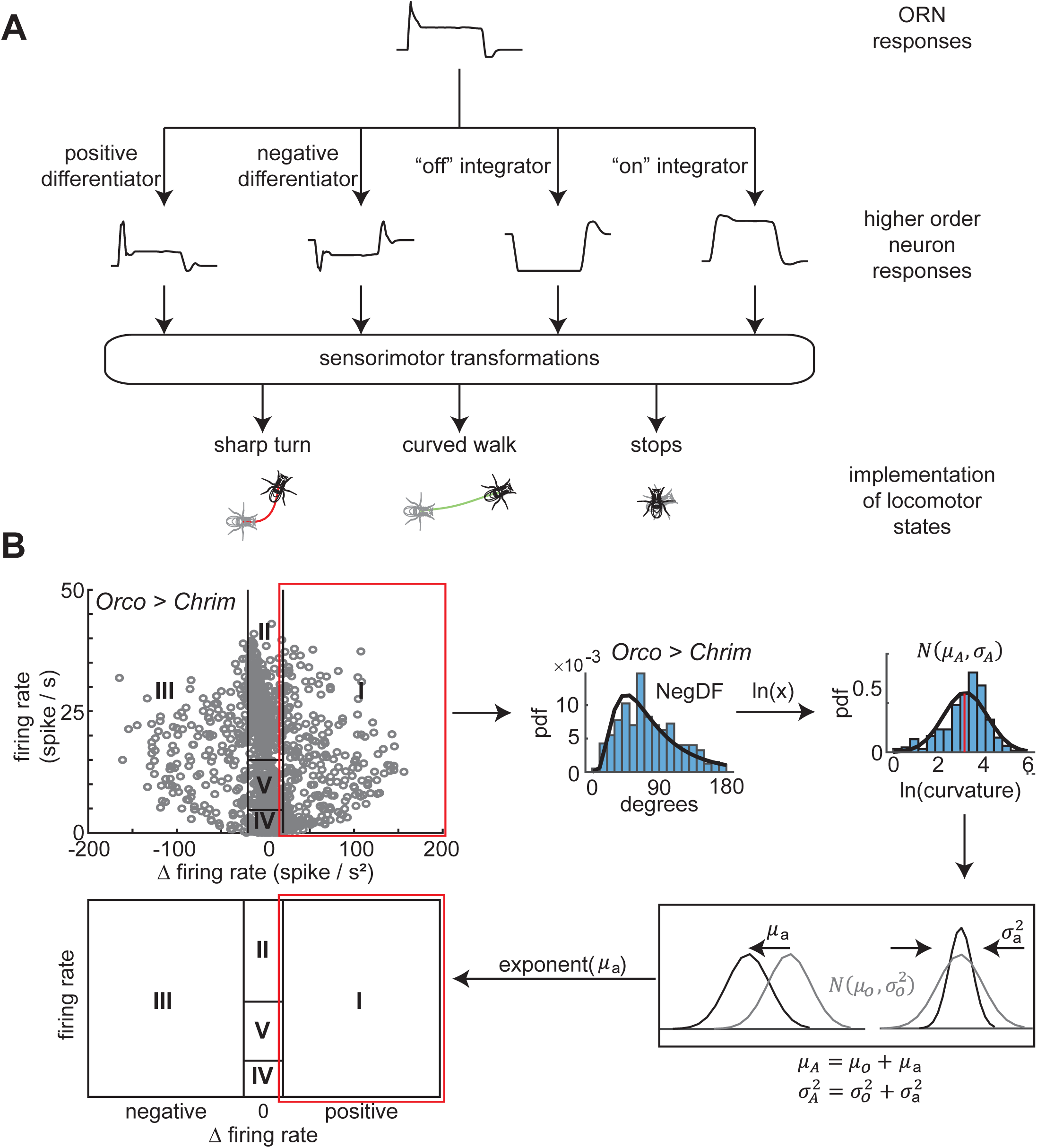
Modeling the effect of a single class (or group) of ORN activity on locomotor kinematics. **A.** Signals from each ORN is carried through parallel channels through higher order neurons that act as positive and negative differentiators and integrators of ORN activity. These channels lead to downstream sensorimotor circuits that modulates the kinematics of different locomotor states. We only show 4 channels for convienience. There can be many more. Our analysis is based on 5 regions. **B.** Locomotor trajectories can be classified into 5 broad regions based on firing rate and change in firing rate. The distribution of locomotor kinematics (sharp turn curvature for *Orco>Chrim* is shown) follows a gaussian distribution after taking a natural log transform. The effect of ORN activity is modeled by the shift in the mean of this distribution as compared to baseline (the distribution prior to first entry) and either increasing or decreasing the variance in the distribution as compared to baseline. The mean shift transformed back into the kinematics space can then be summarized in a schematic reflecting each of the five broad ORN activity regions.

**Figure 6S2.**
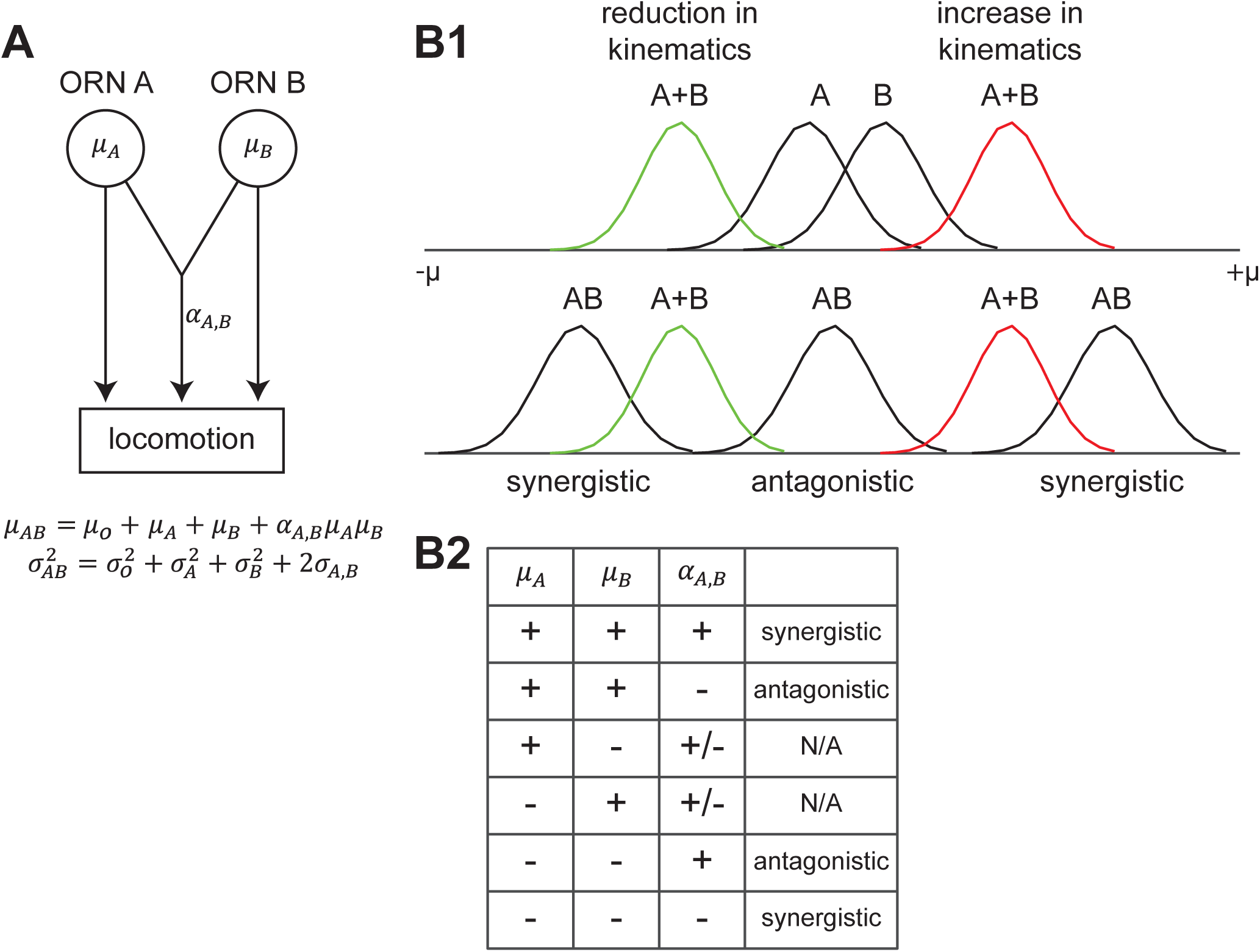
Rules of combination for two sets of ORNs. **A.** Schematic of individual ORN effect on locomotion and an combinatorial effect on locomotion. **B1.** Under separate pathways, the net effect on locomotion when coactivating two sets of ORNs is the sum of each set of ORN. ORNs interact synergistically when ORNs that cause an increase in locomotor kinematics shows an higher increase in kinematics when co-activated than the sum of the individual effects. The reverse is true when ORNs decrease the locomotor kinematics. Meanwhile, if the net effect on locomotion is less than the sum of it’s parts, then the effect is antagonistic. **B2.** Whether any 2 sets of ORNs interact in a synergistic or antagonistic manner is dependent on the sign of the contribution of each individual set of ORN and the interacting term between them.

**Figure 6S3.**
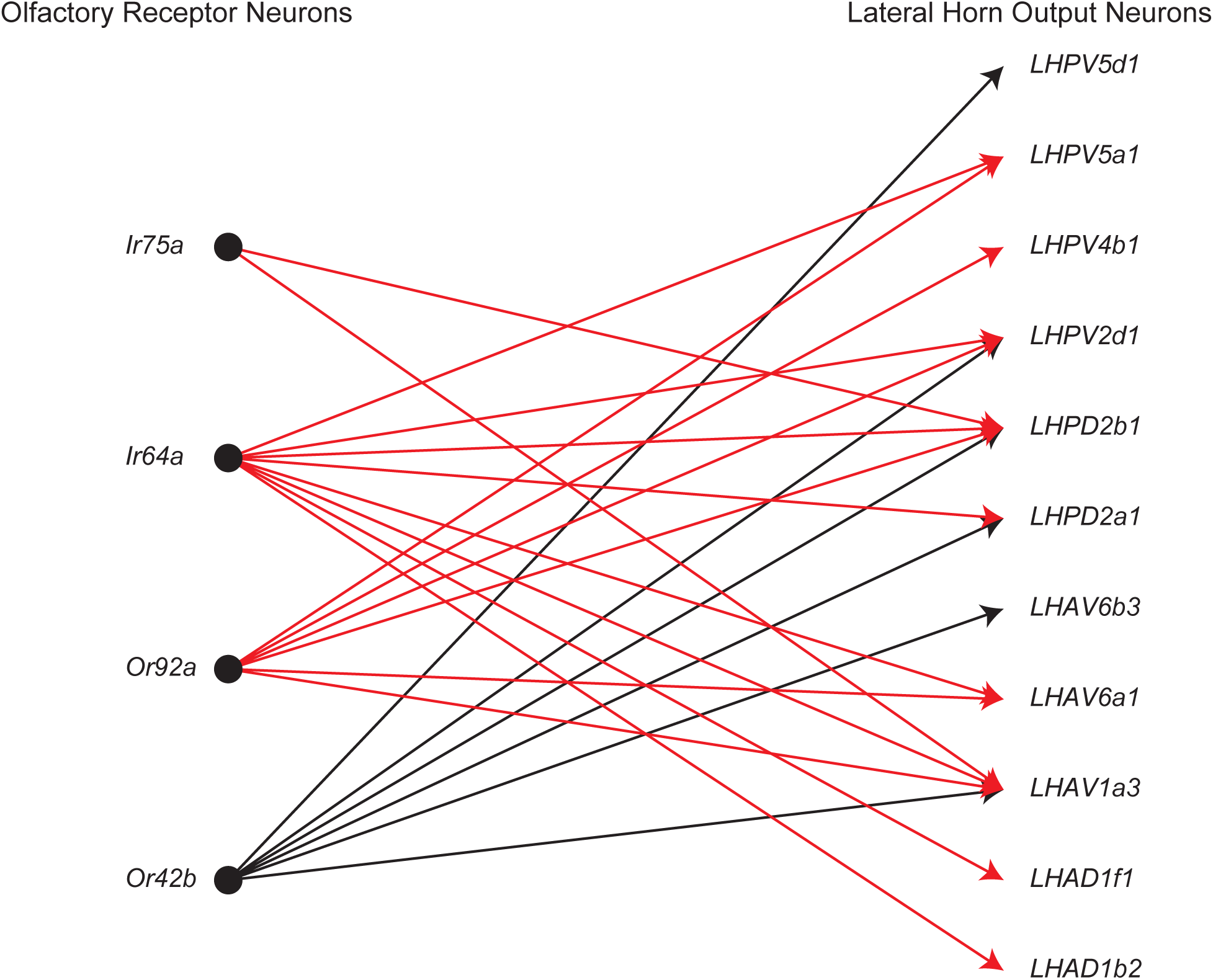
Convergent and divergent integration of olfactory information in lateral horn output neurons (LHON). Each arrow indicates that a direct (via uniglomerular projection neurons) or second order connection (via multiglomerular projection neurons) exists between an ORN class used in this study and the respective LHON class.

**Figure 7S1.**
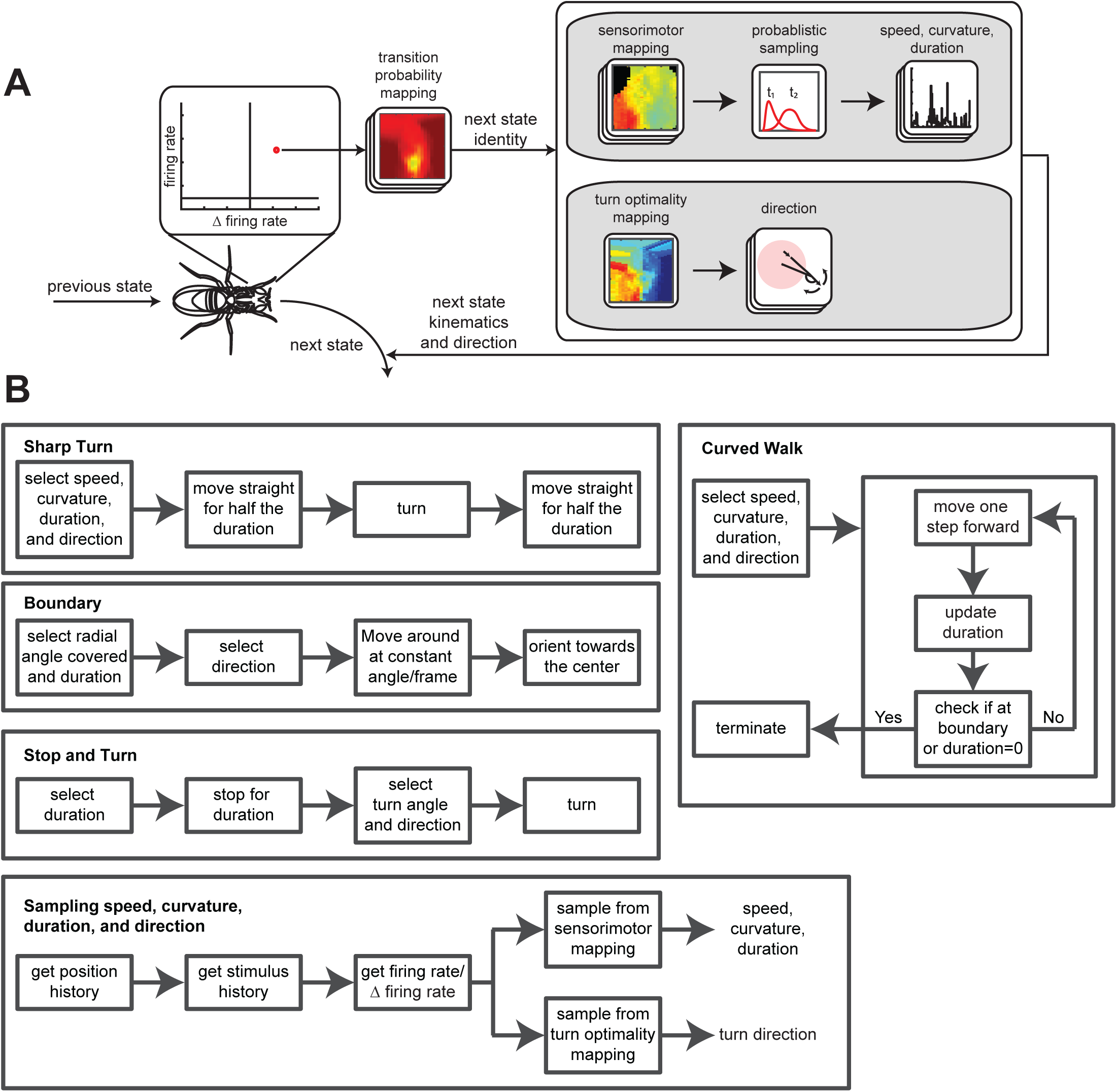
Details of the agent based model. **A.** Synthetic flies can be in one of four states: sharp turn, curved walk, boundary, and stops. During state transitions, flies experience some ORN firing rate and change in firing rate activity. They will transition to a new state based on the ORN activity. Based on the ORN activity and time since first stimulus encounter, they will sample from an lognormal distribution for the new state’s speed, curvature, and duration. Synthetic flies will also choose a direction based on the turn optimality mapping based on the ORN activity and time since first stimulus. **B.** Pipeline of steps synthetic flies take during each of the four states. Like directed runs, a sharp turn will transition into a boundary state if the fly reaches the boundary prior to the end of the sharp turn instance. The fly leaves the boundary by orienting towards the center of the arena at an angular offset of +/- 10 degrees. Adapted from “Mechanisms underlying attraction to odors in walking Drosophila.” by Tao, L., Ozarkar, S., & Bhandawat, V. 2020, PLOS Computational Biology, 16(3), e1007718.

**Figure 7S2.**
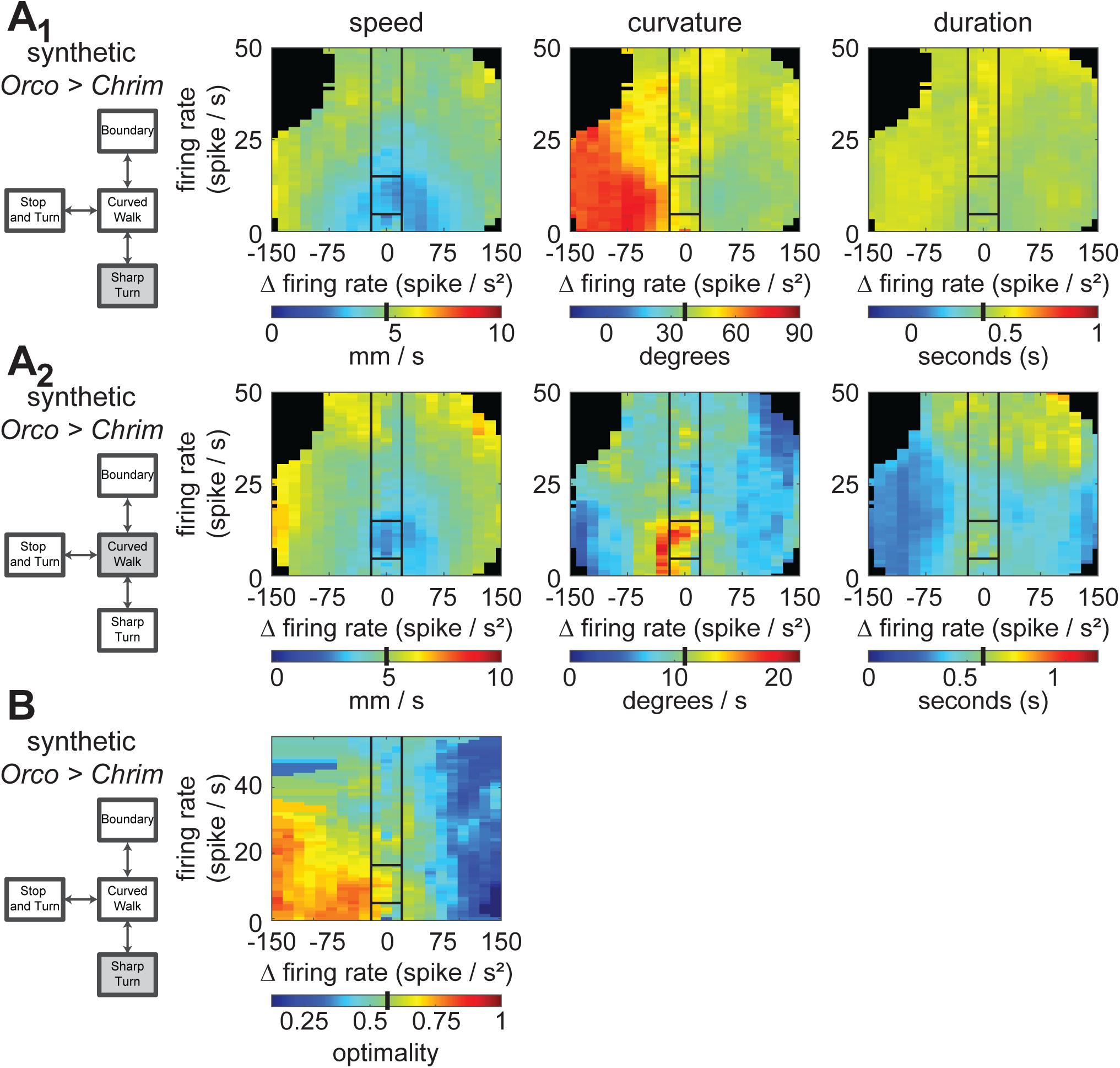
An agent based model of flies based on locomotor kinematics and turn optimality can preserves the sensorimotor mappings of empirical flies. A. Sensorimotor mapping of the effect of ORN activity on Sharp Turn (A1) and Curved Walk (A2) speed, curvature, and duration for synthetic flies. B. Turn optimality (defined as the percentage of optimal turns) for synthetic flies relative to baseline. Black lines separate out each sensorimotor mapping into 5 broad regions based on firing rate and change in firing rate. Baseline turn optimality is shown as a black bar in the color- map.

**Figure 7S3.**
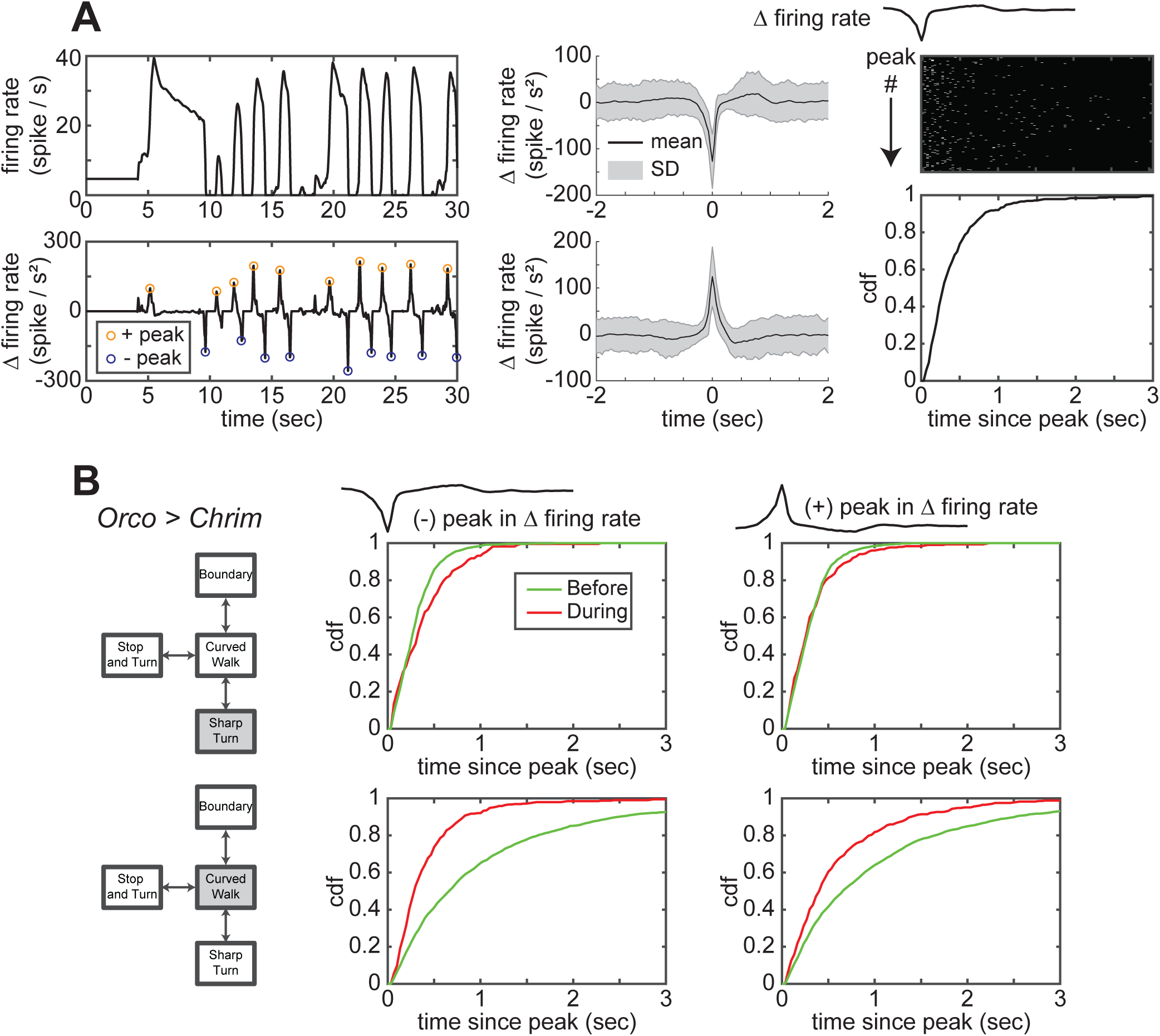
Large change in ORN firing rate causes flies to transition from a curved walk to sharp turn. **A. Left:** Sample 30 second firing rate and change in firing rate of an Orco>Chrimson fly. Circle show positive (orange) and negative (blue) peaks in ORN activity. **Center:** Mean and standard deviation of positive and negative peaks aligned by time of peak in change in firing rate. **Right:** Raster of time after negative peak in change in firing rate when the first transition out of a curved walk state occurs. Each row represents an instance where the fly experiences a negative peak in change in firing rate when in the curved walk state. These transition times are summarized using the cumulative distribution function (cdf). **A.** CDF for sharp turns (top row) and curved walks (bottom row) time to state transition since negative (middle column) and positive (right column) peak in change in firing rate. Before CDFs are constructed by randomly sampling time points to serve as “peaks in change in firing rate” that belong to curved walk and sharp turn tracks before first entry.

**Figure 7S4.**
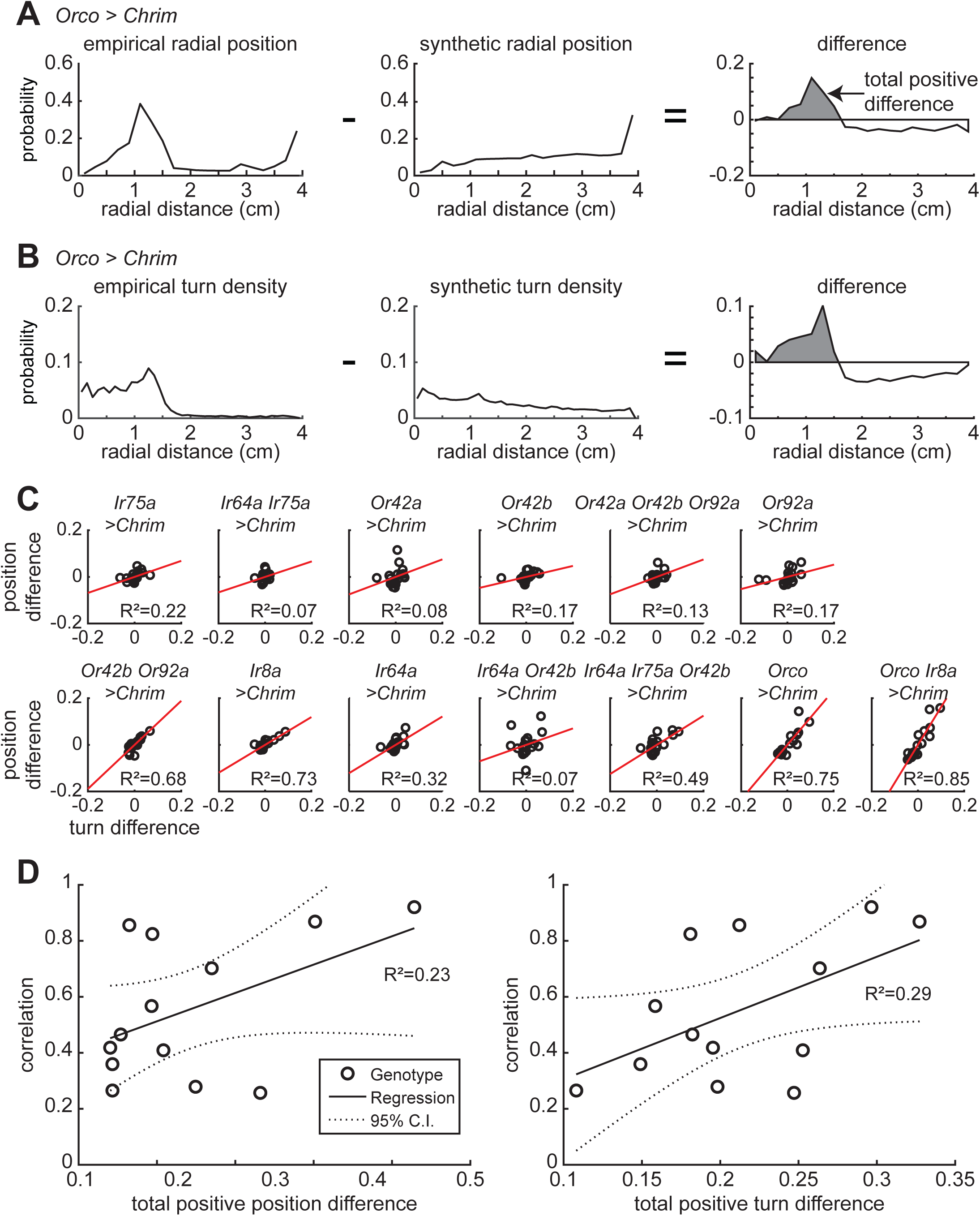
Radial occupancy differences between empirical and synthetic flies is correlated with turn density differences. **A.** Difference in radial occupancy between *Orco > Chrimson* empirical flies (left) and synthetic flies (middle). The total positive difference is calculated represents the amount of change between the empirical and synthetic. **B.** Same as **A**, but for turn density. **C.** Scatter plots of the difference in turn density against difference in radial occupancy for each genotype listed in the same order as the main Figure 7. **D.** The difference in radial occupancy is highly correlated with the difference in turn density for genotypes where there is a large amount of difference in either radial occupancy (left) or turn density (right) between the empirical and synthetic.

